# Functional brain network community structure in childhood: Unfinished territories and fuzzy boundaries

**DOI:** 10.1101/2021.01.21.427677

**Authors:** Ursula A. Tooley, Danielle S. Bassett, Allyson P. Mackey

**Affiliations:** Neuroscience Graduate Group, Perelman School of Medicine, University of Pennsylvania, Pennsylvania, PA 19104, USA; Department of Psychology, School of Arts and Sciences, University of Pennsylvania, Pennsylvania, PA 19104, USA; Department of Bioengineering, School of Engineering and Applied Sciences, University of Pennsylvania, Pennsylvania, PA 19104, USA; Department of Electrical & Systems Engineering, School of Engineering and Applied Sciences, University of Pennsylvania, Pennsylvania, PA 19104, USA; Department of Physics & Astronomy, School of Arts and Sciences, University of Pennsylvania, Pennsylvania, PA 19104, USA; Department of Neurology, Perelman School of Medicine, University of Pennsylvania, Pennsylvania, PA 19104, USA; Department of Psychiatry, Perelman School of Medicine, University of Pennsylvania, Pennsylvania, PA 19104, USA; Santa Fe Institute, Santa Fe, NM 87501 USA

**Keywords:** development, community structure, networks, graph theory, network neuroscience

## Abstract

Adult cortex is organized into distributed functional communities. Yet, little is known about community architecture of children’s brains. Here, we uncovered the community structure of cortex in childhood using fMRI data from 670 children aged 9-11 years from the Adolescent Brain and Cognitive Development study. Children showed similar community structure to adults in early-developing sensory and motor communities, but differences emerged in transmodal areas. Children have more cortical territory in the limbic community, which is involved in emotion processing, than adults. Regions of association cortex interact more flexibly across communities, creating uncertainty for the model-based assignment algorithm, and perhaps reflecting cortical boundaries that are not yet solidified. Uncertainty was highest for cingulo-opercular areas involved in flexible deployment of cognitive control. Collectively, our findings suggest that community boundaries are not solidified by middle childhood, an instability that provides important context for children’s thoughts and behaviors.

## INTRODUCTION

The human cortex is made up of distributed functional communities. Each community is a set of preferentially interacting regions that can be reliably detected across individuals^1–3^. Communities follow a gradient from uni-modal to heteromodal^4^, which aligns with a hierarchical gradient of intrinsic timescales, from fast to slow^5^. Communities comprised of higher-order association areas are expanded in humans compared to other primates^6^, have high expression of genes diverging most swiftly from primates in recent human lineage^7^, and show high interindividual variability in adulthood^2,8,9^. Dorsal and ventral attention communities are involved in processing and acting on sensory information^10^. The default community, in contrast, supports internally-constructed representations of information that cannot be sensed directly, such as remembering the past, envisioning the future, and imagining the minds of others^11^. The frontoparietal community flexibly coordinates other communities to meet task demands^12^, and maintains information no longer present in the environment^13^. The clear organization of functional communities and their mapping to cognitive processes begs the question of how these communities are organized during human development.

Humans have the longest childhood of all primates. Prolonged cognitive immaturity is thought to provide a longer window of sensitivity to the environment^14,15^. After the first decade of life, humans have still not reached adult levels of cognitive control or emotion regulation^16,17^. However, 10-year-olds have made considerable progress in their cognitive development: gains with age in skills like working memory, inhibition, and reasoning are much steeper before age 10 than after^18^. In the brain, by age 10, sensory cortex is relatively mature, but association cortex continues to change well into adolescence^19,20^. Cortical thickness in association cortex declines and surface area transitions to an adult-like pattern at age 10, shifting from early childhood expansion to the protracted decrease that continues through adulthood^21,22^. Myelination, as reflected by diffusion measures, is near-complete in occipital and commissural tracts, but frontotemporal-associated tracts show continuing development through adolescence^23,24^. Some evidence from a recent study of 9- and 10-year-olds suggests that community organization resembles that of adults^25^, although their approach only analyses strong connections, which may obscure finer-grained distinctions in still-developing association cortex. From these studies, one might think that in middle childhood children’s brains are organized more or less like adults. However, it is also true that adolescence involves major cognitive and neural reorganization^26,27^. Accordingly, one might also think that in middle childhood children’s brains are not organized like adults. Which is true?

To probe the organization of children’s brains in middle childhood, we turn to tools from network science, formalizing the brain as a collection of nodes and edges^30^. Network analyses of child brain development have shown increased segregation and integration with age, resulting in the eventual efficient small-world architecture of adult networks^31–35^. However, these studies have largely focused on developmental processes, or employed adult group-level communities to probe changes in network architecture (e.g., Ref.^36^). During adolescence, age is associated with shifts in the boundaries of functional communities^27^, suggesting that the cortical patterning of community boundaries might be quite different in childhood than in adulthood. To our knowledge, however, few studies have explicitly compared cortical patterns of functional communities in childhood with those found in adults. Further, it is an open question whether group-level community assignments might map to individual children more poorly than in adults, because of greater interindividual variability in children. Despite this gap, many developmental studies have applied adult group-level communities, which may lead to incorrect conclusions^37,38^, either because children’s brains are not like adults, or because they are more different from each other^39^.

Here, we sought to test whether the patterns and contours of children’s group-level cortical functional communities resemble those previously found in adults or whether they differ, as we predicted, primarily in higher-order association areas. We first employed a widely-used data-driven approach to cluster points on the gray matter surface based on their patterns of connectivity to the rest of the brain. The clustering approach was developed by Yeo *et al*.^28^ in their influential adult partition. The resulting developmental partition raised further questions about the interactions between communities: to probe these interactions and the reliability of community assignments, we turned to the weighted stochastic block model (WSBM), a generative model-based approach developed by Aicher *et al*.^40^. The WSBM explicitly models interactions both within and between communities, attempting to partition communities such that nodes with similar patterns of connectivity are grouped together. The WSBM also has the biologically-motivated assumption of fewer organizing principles, and thus may align better with large-scale brain organization, and provides a greater flexibility and sensitivity to detect a diverse set of network architectures^41,42^. As many developmental processes, such as synaptic pruning and myelination, are still occurring in middle childhood, we hypothesized that connectivity might still be undifferentiated in higher-order communities, thus in each approach, we examined regions of high and low certainty in community assignment. Finally, to investigate the variability we observed in transmodal association cortex in middle childhood, we chose a task with strong patterns of activation and deactivation in association cortex communities, the n-back task, to test the relationship between children’s community topography and their patterns of task activity.

## RESULTS

### Children show limbic community expansion and greater integration of somatomotor and language communities

We first examined how cortical patterns of functional communities differed in middle childhood from patterns previously found in adults. We used a well-established community detection algorithm, applied to adults in Ref.^28^, to create a group-level developmental partition from vertex connectivity profiles (Figure 1). We investigated the stability of community partitions across different numbers of communities with a resampling approach to determine whether *k*=7 was a reasonable choice. Local minima in instability indicate the number of communities that can stably estimated using the clustering algorithm; we observe a marked increase in instability after *k* = 7 (Figure 1b). Though our primary goal was ease of comparison to the adult 7-community partition in Ref.^28^, our findings of a local minimum at *k*=7, as was found in an adult sample^28^, suggests that 7 communities is an appropriate starting point for partitioning cortex in children.

**FIG. 1.**
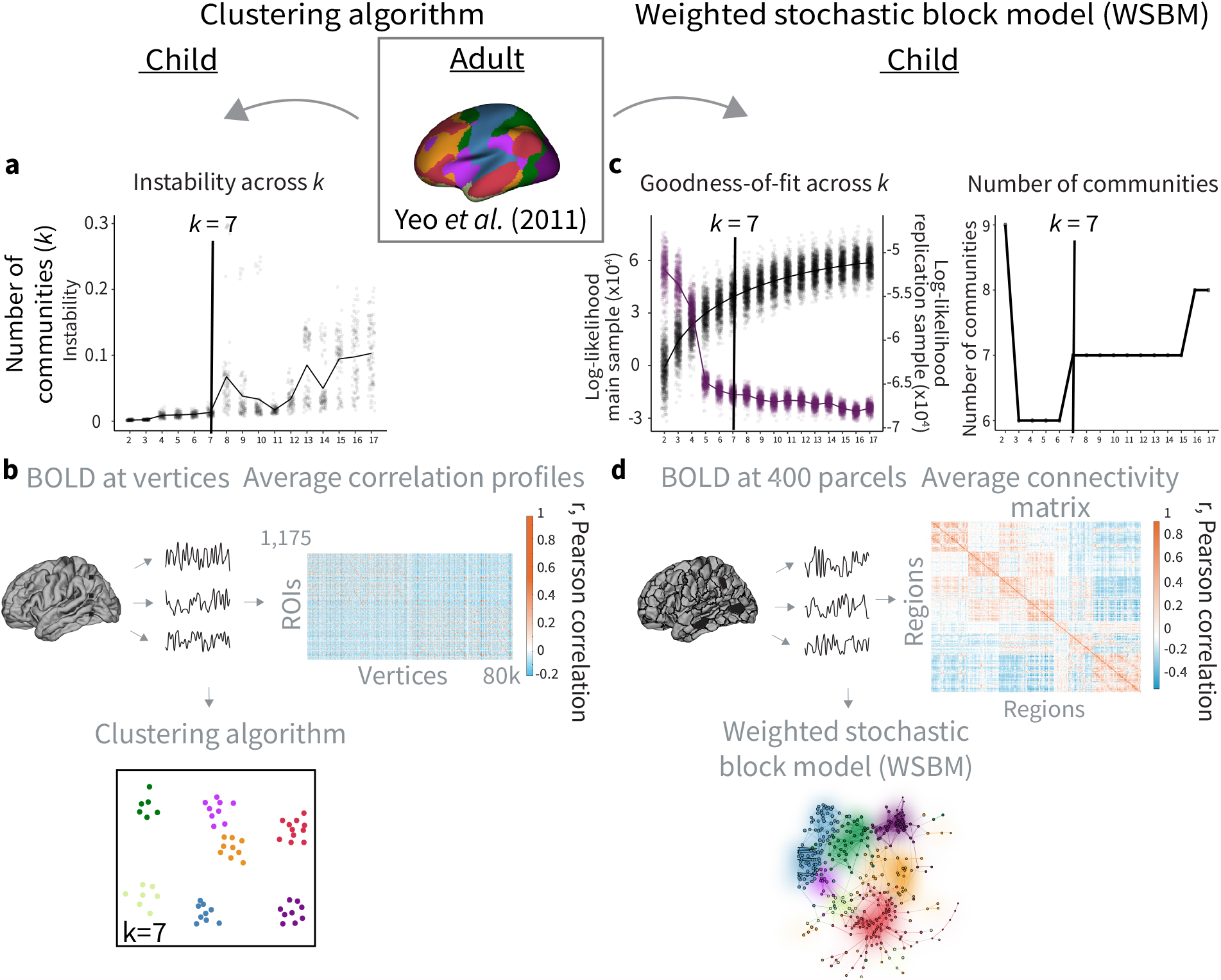
Overview of methods. The adult partition of cortical regions into 7 communities^28^ (inset) was compared to two developmental partitions: one generated using a data-driven clustering approach and one generated using a model-based WSBM approach. **a**, We investigated the stability of different numbers of communities (*k*) using a resampling approach with the clustering algorithm (see *STAR Methods*). With increasing number of estimated communities, we observe less stability, which is expected as the number of estimated communities enlarges the solution space of the clustering problem. Local minima indicate the number of communities that can stably estimated using the clustering algorithm; we observe a marked increase in instability after *k* = 7 (black line). **b**, Schematic of pipeline for generating the clustering partition. Vertex-wise surface data was extracted and correlation profiles across 1,175 equally-spaced regions of interest (ROIs) were calculated for each vertex (average correlation profiles depicted). These profiles were then used as input to the clustering algorithm, which attempts to cluster vertices into *k* = 7 communities. **c**, We investigated the goodness-of-fit (log-likelihood) of different numbers of communities (*k*) across our main (black) and replication (purple) datasets using the WSBM, finding a noted decrease in goodness-of-fit in the replication dataset around *k*= 5 − 6 (left panel). We observed a distinct plateau in the number of communities detected at the group level at *k* = 7 (right panel). **d**, Schematic of pipeline for generating the WSBM partition. Regional timeseries were extracted using a 400-region parcellation^29^, and correlations between regional timeseries were represented as a network (average adjacency matrix depicted). These networks were then used as input to the WSBM, which attempts to group parcels into *k =* 7 communities.

The resulting child clustering partition with 7 communities is shown in Figure 2b. In Figure 2c, we show the allocation of cortex to communities in the clustering partition and a comparison to the adult partition. In the clustering partition, more total surface area was assigned to the limbic and visual communities than in the adult partition, and less to the default and frontoparietal communities than in the adult partition. This observation is suggestive of a relative expansion of limbic and visual territory, and contraction of default and frontoparietal territory, in children relative to adults.

**FIG. 2.**
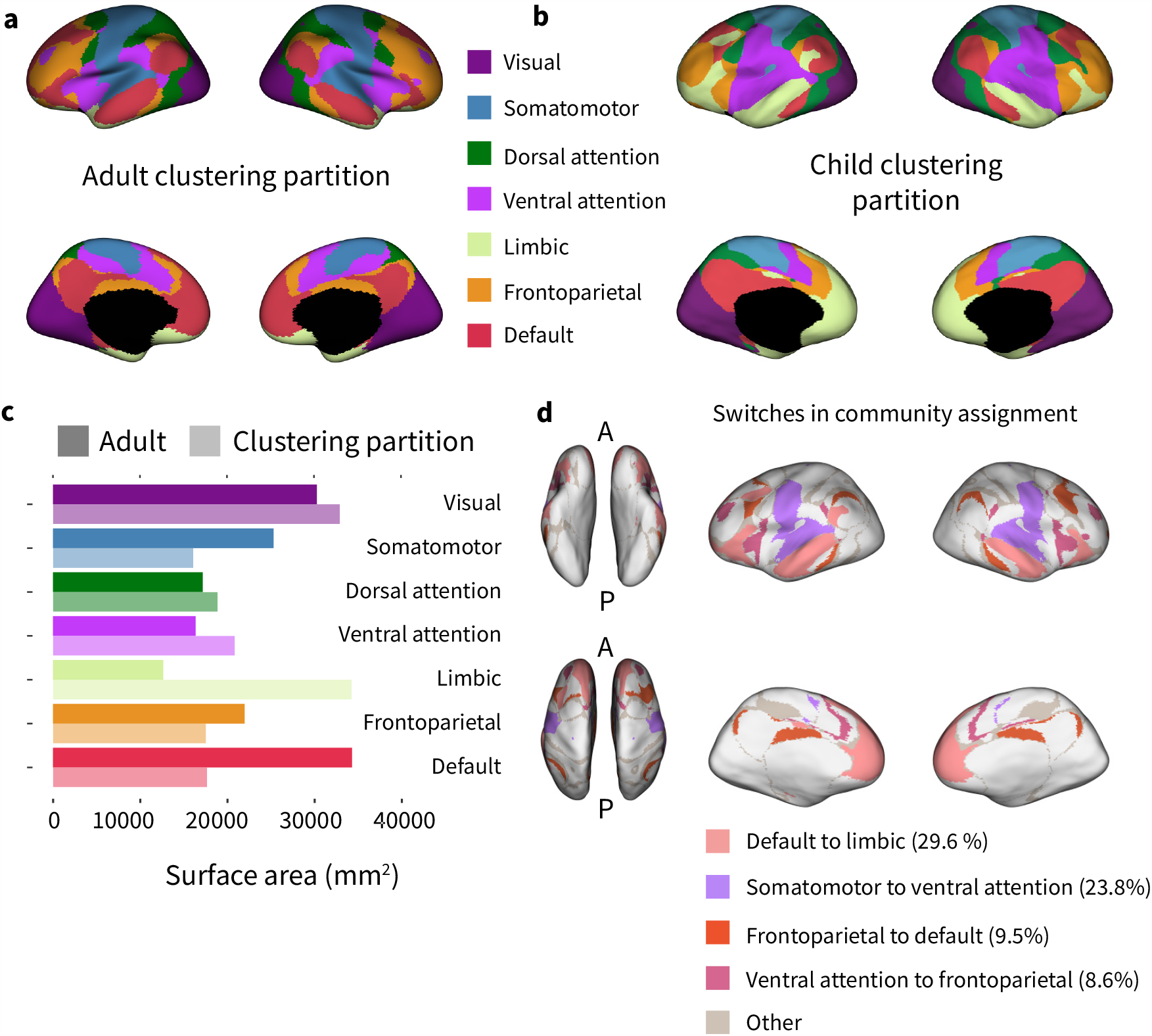
Overview of partitions generated using a data-driven clustering approach. **a**, The partition of cortical regions into 7 communities estimated by Ref.^28^ by applying a clustering approach to adult neuroimaging data. **b**, A partition estimated from developmental data using the same clustering approach. Note the overall similarity between the child partition and the adult partition (normalized mutual information (NMI) = 0.64, *p* < 0.001, permutation test). **c**, Surface area assigned to each community in the two partitions. **d**, Areas that were assigned to different communities in the adult partition and the child partition. Switches in community assignment from the adult partition to the child partition are shown in color.

To further probe the differences between the contours of children’s functional communities and those of adults, we investigated measures of partition similarity and the specific brain regions that showed differences in community assignment. The clustering partition is significantly more similar to the adult partition than expected by chance: the normalized mutual information of the two partitions is 0.64 (*p* < 0.001, permutation test), and the normalized information distance^43^ is 0.36 (*p* < 0.001, permutation test). Further, 39.53 % of vertices (ignoring the medial wall) have a different assignment in the clustering partition than in the adult partition. Of these vertices, the majority of switches were in assignment of (i) adult default system regions to the limbic community in children (29.63 %), and of (ii) adult somato-motor regions to the ventral attention community in children (23.8 %) (Figure 2d). 9.46 % of switches were from the frontoparietal to the default community, and 8.63 % were from the ventral attention to the frontoparietal community. The ventral regions of the precentral (primary motor cortex) and postcentral gyri (primary somatosensory cortex), regions that typically encode the face^44^, are clustered with the ventral attention community, a fractionation that was previously observed in the 17-community partition in adults in Yeo *et al*.^28^.

### Uncertainty in community assignment is high in transmodal regions

When connectivity is distributed evenly in a similar pattern across communities, the assignment of regions to communities will be uncertain, whereas when connectivity is clearly segregated into differentiated patterns, the assignment of regions to communities will be certain. Here we sought to understand where connectivity may not yet be clearly segregated in the child brain, and we therefore calculated the certainty in the assignment of regions to communities. We employed the confidence measure used in Ref.^28^ to index certainty in community assignment across vertices (see Methods); higher values of confidence are indicative of higher certainty in community assignment. In the adult sample examined by Ref.^28^, areas of low confidence fall primarily along borders between communities, and sometimes indicate where communities could be fractionated in a higher-resolution partition (Figure 2a). Similarly, areas of low confidence also fall along boundaries between communities in the clustering partition (Figure 2b), but there are additional regions in the clustering partition that show low confidence in assignment in the posterior cingulate, precuneus, and inferior parietal lobule (circled in Figure 3a and 3b). Despite lower confidence, the precuneus and posterior cingulate area maintain similar community assignment across both the adult partition and the two developmental partitions (Figure 2b and 4a-c), varying only in their spatial extent. Similarly to adults, areas of low confidence in children seem to primarily indicate fuzzy delineations between communities, but it is difficult to ascertain visually whether the certainty along boundaries of higher-order communities is lower than the certainty along boundaries of primary sensory areas. For that, we turn to an investigation at the level of functional systems.

**FIG. 3.**
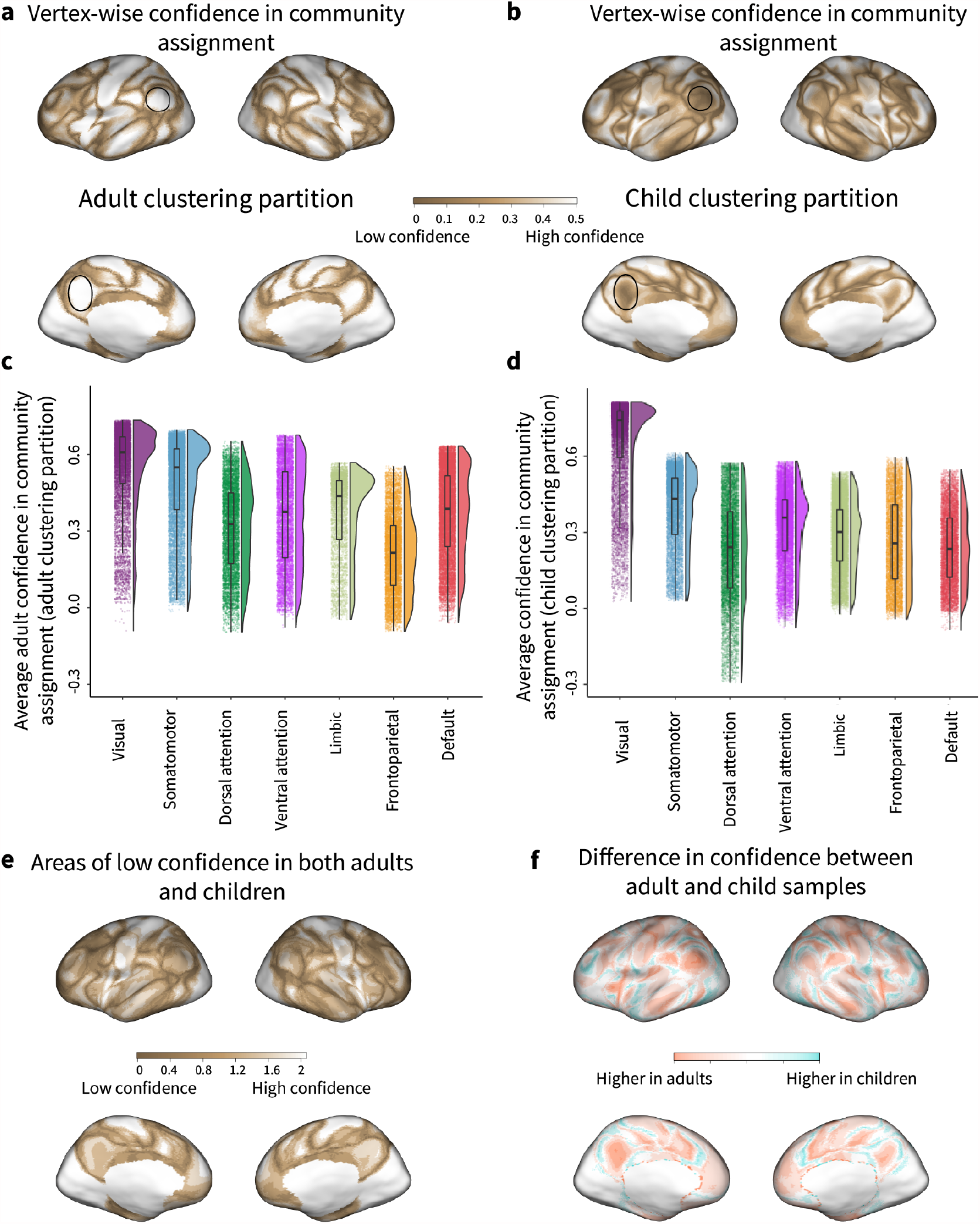
Confidence in community assignment using a data-driven clustering approach. **a**, Confidence maps estimated from an adult sample^28^ using the silhouette method. This method measures the similarity of a given vertex’s timeseries to that of other vertices assigned to the same community, compared to the next most similar community (see Methods). For the purposes of visualization, negative silhouette values were set to 0. **b**, Confidence maps estimated from the developmental sample using the same method. For the purposes of visualization, negative silhouette values were set to 0. **c**, Average confidence in the adult sample within each of the systems in the adult partition. **d**, Average confidence in the developmental sample within each of the communities in the child clustering partition. Note the higher confidence in the visual community and relatively lower confidence in higher-order association communities, particularly the dorsal attention and default communities. **e**, Summed confidence in community assignments from adult and developmental samples. Dark brown indicates areas of low confidence in both adult and developmental samples. **f**, Difference in confidence maps between adult and developmental samples. Red indicates higher confidence in the adult sample, while blue indicates higher confidence in the child sample; these differences should be interpreted with caution, as there are other differences between the adult data and the developmental data used here.

We asked whether fuzzy delineations of boundaries are distributed broadly across association cortex in both childhood and adulthood, or whether children show areas of undifferentiated connectivity primarily within the limbic and somatomotor systems that are assigned differently than in adults. We quantified this by calculating the median confidence within each community in both the adult and child samples (Figure 3c,d). In the adult sample examined in Ref.^28^, confidence was highest in vertices assigned to the visual and somatomotor systems and slightly lower in vertices assigned to higher-order association systems (Figure 3c, *H* (6) = 21179.38, *p* < 2 × 10^−16^). In particular, there was lowest confidence in assignment in regions in the frontoparietal (all pairwise comparisons *p* < 0.01, Bonferroni corrected) and dorsal attention systems (all pairwise comparisons significant except ventral attention, *p* < 0.01, Bonferroni corrected). In our developmental sample, confidence was again highest in vertices assigned to the visual community, with significantly lower confidence in vertices assigned to other communities (Figure 3d, *H* (6) = 25933.24, *p* < 2 × 10^−16^). There was lowest confidence in assignment in regions assigned to higher-order association communities, in particular, the dorsal attention (all pairwise comparisons *p* < 0.01, Bonferroni corrected), default (all pairwise comparisons significant except dorsal attention *p* < 0.01, Bonferroni corrected), and frontoparietal systems (all pairwise comparisons significant, *p* < 0.01, Bonferroni corrected). These results are robust to using the community assignments from the adult clustering partition instead of the child clustering partition (see Figure S1a, *H* (6) = 28310.69, *p* < 2 × 10^−16^), finding again that confidence was highest in vertices assigned to the visual system, and lowest in frontoparietal (all pairwise comparisons *p* < 0.01, Bonferroni corrected), dorsal attention (all pairwise comparisons *p* < 0.01, Bonferroni corrected), and default (all pairwise comparisons except ventral attention *p* < 0.01, Bonferroni corrected). Note that values of confidence in the developmental sample are overall lower than those of the adult sample used in Ref.^28^, though we do not conduct statistical tests comparing the two, as this difference could be due to other discrepancies between the adult data and the developmental data used here. Our results suggest that in the child brain, connectivity in higher-order association regions, particularly the dorsal attention, default, and frontoparietal communities, is not yet clearly segregated.

We wondered whether areas of undifferentiated connectivity in children were similar or different to those found in adults. We began by comparing the spatial distributions of confidence in the adult and child samples. Summing the confidence maps, we found that areas of low confidence are predominantly in higher-order association cortex in both children and adults (Figure 3e). Visual cortex and the somatomotor strip show relatively high confidence in both samples. Examining the differences in confidence between adult and child samples, we found that overall, children show lower confidence in community assignment (Figure 3f). We did not strongly interpret this difference because it could arise from several distinct differences between the adult data used by Ref.^28^ and the child data used here. Note that visual areas show no differences in confidence between adult and child samples, as they are relatively high confidence in both adults and children. Overall, these results above suggest that children’s cortical patterns of connectivity may be less differentiated than those of the adult brain.

When examining the distribution of certainty across communities, we found that the dorsal attention community in particular had some regions with very low values of confidence. Investigating these values, we found that these regions were located along the border between the visual and dorsal attention communities (see Supplemental Figure S2). Some of these areas are assigned to the visual community in the adult clustering partition (see negative values in the visual community in Supplemental Figure S1). We observe the assignment of a coherent region in the superior parietal lobule, adjacent to the intraparietal sulcus, to the visual community in the child clustering partition. In the adult partition, a smaller region in this area is assigned to the visual system. This is also consistent with the increased surface area allocated to the visual community in the child clustering partition, compared to in the adult partition (see Figure 2c), suggesting that these areas may be more tightly linked to extrastriate cortex (area MT and anterior MT) in childhood than in adulthood.

### Data-driven community assignments are highly stable across samples

If children’s patterns of brain connectivity are simply more different from each other (*i*.*e*. higher interindividual variability) than adult’s patterns of brain connectivity are, then partitions derived from developmental data might be less stable across samples. Therefore, we employed a replication dataset of children from the Adolescent Brain and Cognitive Development (ABCD) study to assess the reliability and generalizability of our findings. Identical pre-processing and clustering algorithm implementations were used on the replication dataset, drawn from two ABCD sites. Community assignments in the replication clustering partition are highly consistent with those generated using the original dataset (see Figure S4a and S4b). A total of 94.99 % vertices are assigned to the same community across both datasets, with 94.76 % of vertices in the right hemisphere and 95.21 % of vertices in the left hemisphere being assigned to the same community. Normalized mutual information of the two partitions is 0.88, and normalized information distance is 0.12. These results suggest that the cortical patterning of functional communities in middle childhood is stable, and reliably shows differences from adult community organization.

### children show less-solidified higher-order community interactions in association cortex

Using the data-driven approach, we observed that there were stable and reliable patterns of cortical community structure in middle childhood that differ from those established in adults. Notably, we found considerable reapportionment of adult default system regions to the limbic community, as well somatomotor regions to the ventral attention community. This raises the question of the origins of these assignments–is it due to changes in connectivity between these specific systems, or broader differences in patterns of connectivity across all other functional communities? To address this question, we turned to another community detection approach, a generative model-based approach called the weighted stochastic block model (WSBM), that differs from the data-driven approach in several important ways. For one, it explicitly models the interactions between communities. While the data-driven approach models patterns of connectivity and attempts to group regions with similar patterns together into a community, the model-based approach partitions cortex by maximizing the likelihood that each “block” of connections between two communities is internally similar and coherent. Additionally, using WSBM allows us to assess not just the reliability of these cortical patterns of communities across approaches, but also employ a more biologically-motivated method with fewer motivating principles.

We first examined whether cortical patterns of communities estimated using the model-based approach resemble those found in adults or those found using the data-driven approach in children. We used the WSBM to create a group-level developmental partition from average parcel connectivity patterns (Figure 1). To determine whether 7 communities was a reasonable choice when using the model-based approach, we first systematically investigated the goodness-of-fit of the WSBM across different numbers of communities (*k*). When fitting the WSBM to our main dataset, we observe the goodness-of-fit steadily increases with increasing *k* (Figure 1c, left panel, black). However, when calculating the goodness-of-fit of the WSBM partition at a given *k* on our replication dataset, we observe a noted decrease in goodness-of-fit as *k* increases past *k =* 6 (Figure 1c, left panel, purple). Furthermore, we examined the number of communities detected at the group consensus level using the WSBM across different values of *k*, and observe the longest distinct plateau at *k =* 7, indicative of a stable group partition^40,45^ (Figure 1d, right panel).

The resulting WSBM partition with 7 communities is shown in Figure 4c. In Figure 4d, we show the allocation of cortex to communities in the WSBM partition and a comparison to the adult partition. In the WSBM partition, more total surface area was assigned to the default, visual, and somatomotor communities than in the adult partition, and less to the attentional communities than in the adult partition. By visually comparing Figures 4a-c, we observed that the WSBM partition resulted in a more diffuse, scattered pattern of community assignments in prefrontal cortex, suggesting that the variability in assignment seen in the clustering partition in the assignment of adult default and somatomotor regions to limbic and ventral attention communities, respectively, may be broadly distributed across association regions of cortex rather than isolated to those specific systems.

**FIG. 4.**
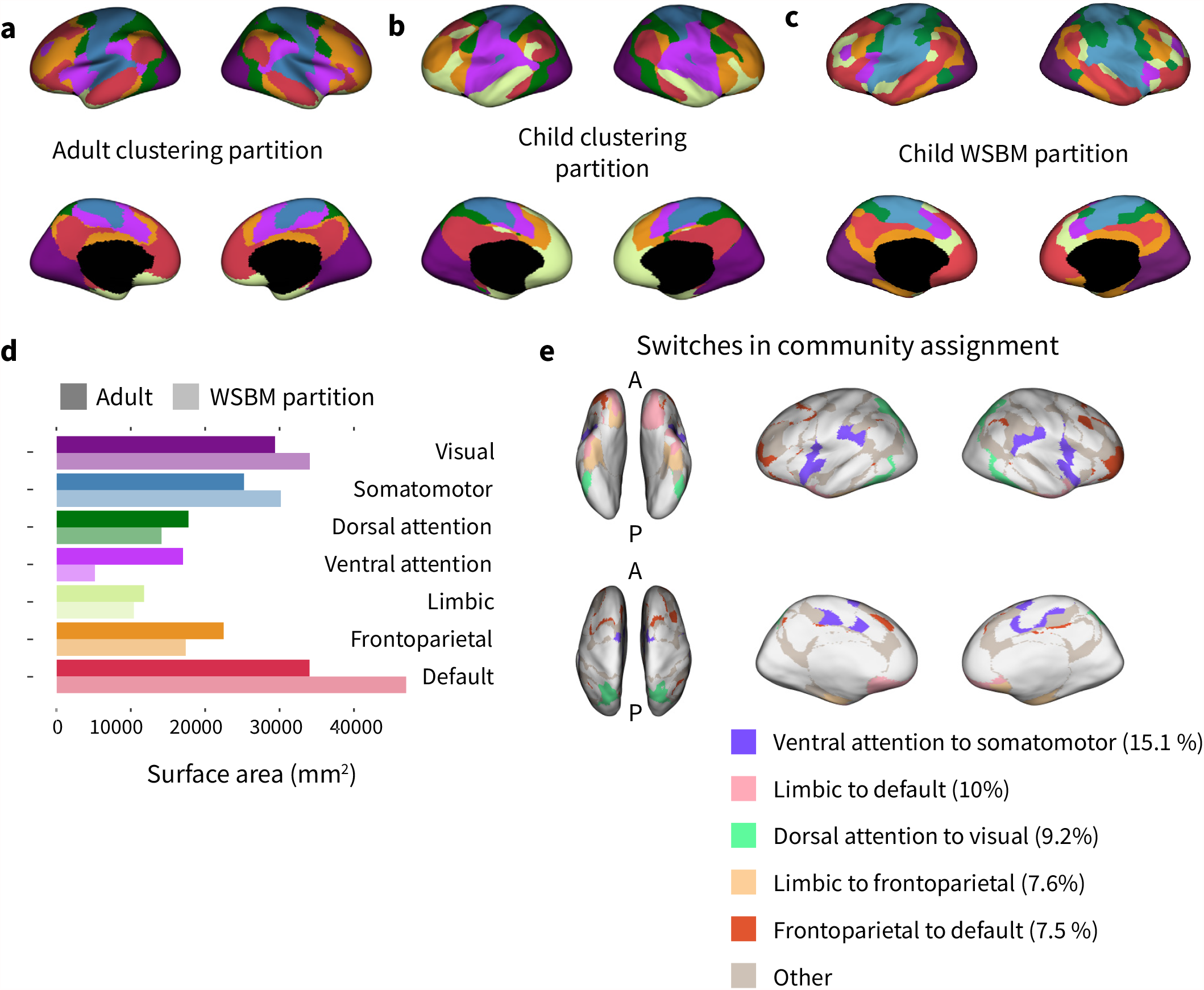
Child partition generated using the model-based WSBM approach. **a**, The 7-community adult partition^28^. Colors in panels a-c correspond to the communities shown in panel d. **b**, A partition estimated from developmental data by applying the clustering approach (see Fig 2b). **c**, A partition estimated from developmental data by applying the WSBM approach. Note the high overall similarity to the adult partition (NMI=0.58, *p* < 0.001, permutation-based testing). **d**, Surface area assigned to each community in the adult partition and WSBM partition. **e**, Areas that were assigned to different communities in the adult partition and the child WSBM partition. Switches in community assignment from the adult partition to the child partition are shown in color.

To probe whether the pattern of functional communities estimated in the WSBM partition differs from that of the adult partition in a consistent fashion, we again examined the specific brain regions that showed differences in community assignment. The WSBM partition is significantly more similar to the adult partition partition than expected by chance: the normalized mutual information of the two partitions is 0.58 (*p* < 0.001, permutation test), and the normalized information distance is 0.44 (*p* < 0.001, permutation test; see Figure 4c). Note that the WSBM partition is more different from the adult clustering partition than is the child clustering partition. Further, 37.23 % of vertices (ignoring the medial wall) have different assignments than in the adult partition. Of these vertices, the majority of switches were in assignment of (i) adult ventral attention system regions to the somatomotor community in children (15.11 %), and of (ii) adult limbic regions to the default community in children (9.98 %) (Figure 4e). 9.18 % of switches were from the dorsal attention to the visual community, 7.6 % were from the limbic to the frontoparietal community, and 7.52 % were from the frontoparietal to the default community. Visual inspection of Figure 4c demonstrates a reversal of the pattern observed in the clustering partition, where face and head areas of the somatomotor community were clustered with the ventral attention community. In the WSBM partition, these regions, which include the insula and parts of auditory cortex, are preferentially grouped with the somatomotor community. The switches in community assignment observed in the WSBM partition somewhat recapitulate the switches in seen in the clustering partition, but suggest that while adult default and ventral attention systems regions do interact differently with the limbic and somatomotor communities (respectively) as seen in the clustering partition, interactions between communities in higher-order association cortex more broadly may be flexible in childhood.

### Uncertainty in assignment is high in regions supporting attentional processes

Next, we sought to examine where interactions between communities might still be quite flexible and undifferentiated, reasoning that the scattered pattern of community assignment in prefrontal areas might be indicative of less-solidified community boundaries in those areas. We employed a measure of variability in community assignment derived from the consensus partitioning algorithm (described in *Methods: Consensus partition algorithm*) to index uncertainty in community assignment. The majority of parcels were assigned to the same community across optimizations of the consensus partitioning algorithm, as we expected, but 48 % of parcels were not consistently assigned across optimizations. We used the proportion of inconsistent community assignments across optimizations to index uncertainty in community assignment. As in the clustering partition, areas of high variability in community assignment are predominantly in higher-order association cortex, found in rostrolateral prefrontal cortex, anterior cingulate, and insula (Figure 5a). These results suggest that in these brain regions, patterns of connectivity are not clearly segregated, and interactions between communities may still be quite flexible. We next turned to investigation at the level of cognitive systems, and asked whether areas of low certainty are distributed broadly across association cortex, or whether they are confined to specific communities. To quantify whether uncertainty in community assignment varied in a systematic manner across the cortex, we calculated the percentage of parcels within each community that were inconsistently assigned (Figure 5b). Variability in assignment was low in regions assigned to primary sensory systems, and varied only to any large extent in higher-order association regions (*H* (6) = 198.34, *p* < 2 ×10^16^). In particular, there was the highest variability in assignment in regions in the ventral attention (all pairwise comparisons, *p* < 0.01, Bonferroni corrected) and limbic (all pairwise comparisons *p* < 0.01, Bonferroni corrected) communities. These results are robust to using the system assignments from the adult clustering partition instead of the child WSBM communities; we find again that the highest variability in assignment was in the ventral attention community (Supplemental Figure S1b, (*H* (6) = 83.68, *p* < 2 ×10^16^), all pairwise comparisons except frontoparietal, *p* < 0.03), though areas defined as part of the frontoparietal system in the adult clustering partition also show high variability in assignment (pairwise comparisons with visual, somatomotor, and default communities, *p* < 0.01, Bonferroni corrected). Taken together with findings from the data-driven approach, our findings suggest that complex undifferentiated connectivity patterns are primarily present in middle childhood in higher-order areas, particularly those supporting attentional processes.

**FIG. 5.**
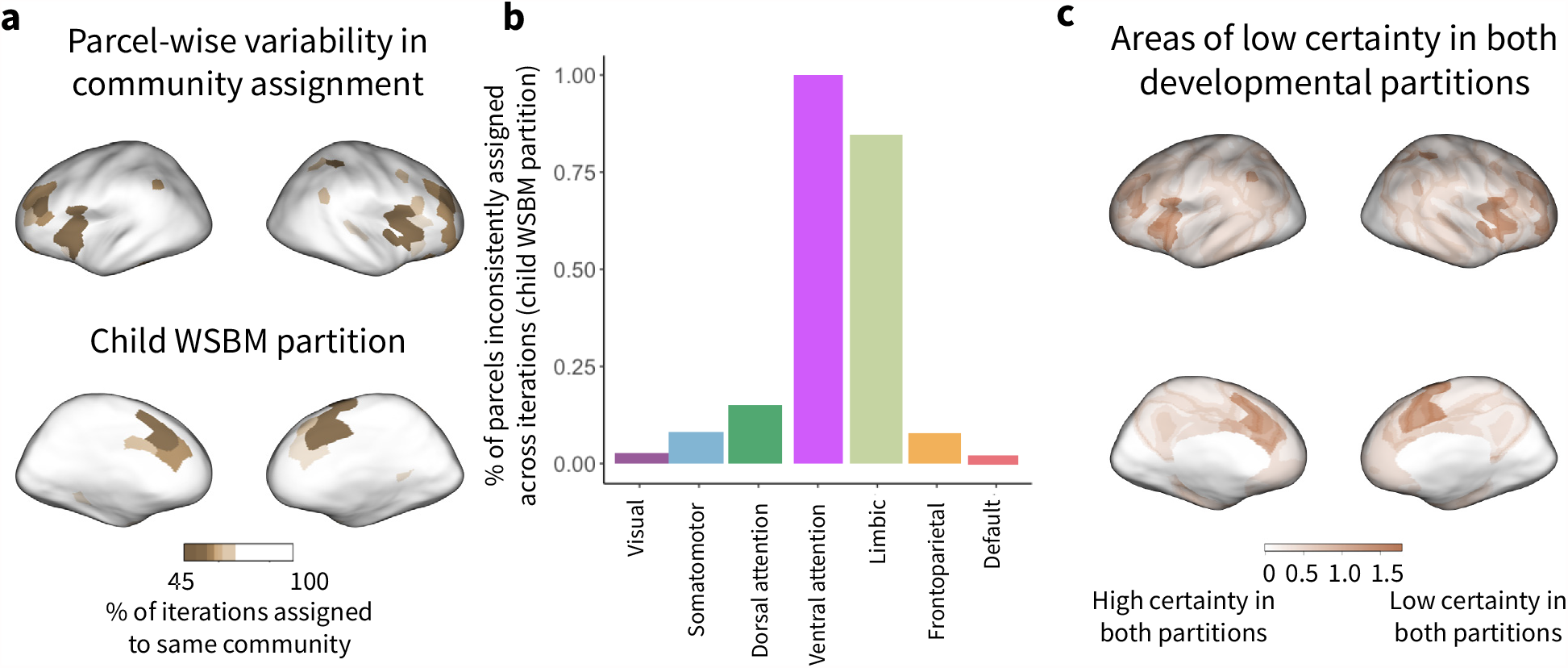
Variability in community assignment using a model-based WSBM approach. **a**, Maps of variability in community assignment estimated from the developmental sample. Parcels that were inconsistently assigned to communities during optimizations of the WSBM consensus partition algorithm are shown in brown. The majority of parcels were consistently assigned to the same community across optimizations. **b**, Variability in community assignment within each of the communities in the child WSBM partition. Note that variability in parcel assignment predominates in the ventral attention and limbic communities. **c**, Areas of low certainty in community assignments across both developmental partitions, regions of low certainty are shown in dark sandstone. Low certainty across approaches is localized to cingulo-opercular regions, namely, the anterior insula and anterior cingulate. Percentage of inconsistent assignment in the WSBM partition is scaled to [0,1] and inverted, then summed with confidence values from the clustering partition scaled to [0,1]. Values close to zero show high certainty in both partitioning approaches.

### Model-based community assignments are moderately stable across samples

To determine the reliability and generalizability of our findings, we repeated our WSBM analyses using a replication dataset of children from ABCD. Identical preprocessing and WSBM implementations were used on the replication dataset drawn from two ABCD sites. Community assignments in the child WSBM partition generated from the replication dataset are somewhat consistent with those generated using the original dataset (see Figure S5a and S5b). A total of 74.62 % of parcels are assigned to the same community across both datasets, with 73.63 % of parcels in the right hemisphere and 75.62 % of parcels in the left hemisphere being assigned to the same community. Normalized mutual information of the two partitions is 0.84, and normalized information distance is 0.17. These results suggest that in middle childhood interactions between functional communities are slightly less stable than the broad cortical patterning of connectivity detected by the data-driven approach. Much of cortex outside prefrontal areas, however, especially primary sensory areas, has clearly-defined stable interactions between communities, which are reliably detectable across samples and across approaches.

### Across both approaches, uncertainty in community assignment is high in cingulo-opercular regions

Finally, we sought to determine where complex undifferentiated connectivity patterns might be present in the child brain, comprising areas where connectivity is not clearly segregated and where interactions between communities remain flexible. Thus, we examined areas of high uncertainty in community assignment across both community detection approaches we employed. We combined the confidence measure from the data-driven community detection approach and the variability measure from the model-based approach to examine the overlap of areas where it was difficult to assign a community identity. The percentage of inconsistent assignments in the WSBM partition was scaled to [0,1] and inverted, and confidence values from the clustering partition were scaled to [0,1]; then the two estimates were summed (Figure 5c). Insular and anterior cingulate areas show low certainty in both approaches, suggesting that during middle childhood, these regions may still be flexibly associated with several functional communities.

We found a similar spatial distribution of regions of low confidence across approaches when using the silhouette measure to index uncertainty in community assignment in the WSBM partition. We previously used the silhouette measure when examining uncertainty in assignment in the child clustering partition. Higher values of the silhouette measure are indicative of higher confidence in community assignment, while lower values of confidence are indicative of lower confidence. Regions of low confidence in the WSBM partition were primarily located in the insula, lateral prefrontal areas, and cingulate (Supplemental Figure S6a). Regions of low confidence using the silhouette measure covered more territory than regions of high variability in community assignment using the measure derived from optimizations of the consensus community algorithm (compare Figure 5a to Supplemental Figure S6a). Insula and anterior cingulate areas again showed low confidence in assignment across both partitioning approaches, though additional regions in the parahippocampal and entorhinal cortex also showed low confidence across partitioning approaches when using the silhouette measure of confidence (Supplemental Figure S7b). We qualitatively observed that posterior regions seemed to have more consistently similar community assignments across partitions. To quantify this observation, we calculated for each vertex the number of different communities it was assigned to across the three partitions. We observed that anterior higher-order association regions tended to vary in assignment more across the three partitions (Supplemental Figure S7b), suggesting that these transmodal areas do not yet have solidified community identities in middle childhood.

### The clustering partition and the adult partition capture functional organization during task performance well

Functional communities have been shown to comprise brain regions that are co-activated during performance of specific cognitive tasks^2,3,46^. In the ABCD study, three tasks were collected: an inhibitory control task (stop-signal), a reward processing task (monetary incentive delay), and a working memory (n-back) task^47^. The n-back task effectively localizes both the frontoparietal community (activity greater than baseline), and the default community (activity less than baseline)^47^. No tasks specifically localized sensory, motor, or attention communities.

#### Frontoparietal community

If the frontoparietal community simply comprises different spatial territory in middle childhood than in adulthood, we would expect to see the differences in the topography of the frontoparietal community found in our developmental partitions reflected in the spatial extent of activation during the n-back task. To examine whether this indeed is the case, we thresholded the task activation maps to retain only the highest 20 % of the contrast weights (Figure 6b), and compared these to the frontoparietal communities in each partition. Task activation in the 2-back versus 0-back contrast of the n-back is shown in Figure 6a. The clustering partition and the adult partition show equally good correspondence to task activation in the n-back task (clustering partition Sørensen-Dice coefficient = 0.8052*±*0.001(0.8025−0.808), adult partition Sørensen-Dice coefficient = 0.8078*±*0.001(0.8051−0.8105)), with the WSBM partition doing worse in comparison (Sørensen-Dice coefficient = 0.7445 ± 0.001(0.7416 − 0.7473)). This set of findings suggests that both the clustering partition and adult partition capture community structure that corresponds well to task activity.

**FIG. 6.**
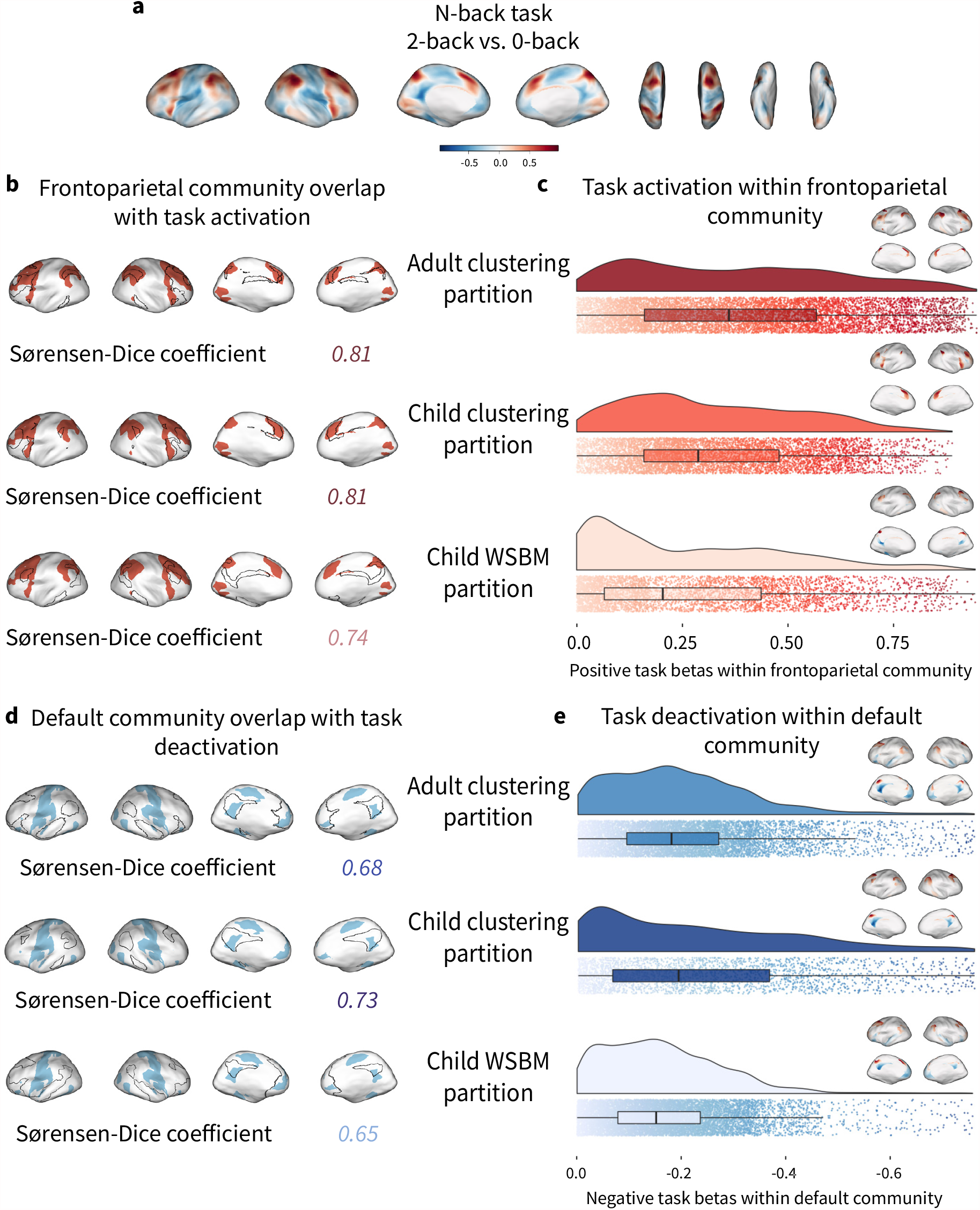
Overlap between task activation and communities in the estimated partitions. **a**, Contrast of 2-back versus 0-back in the emotional n-back task, controlling for age, sex, scanner, and performance. **b**, Overlap of highest 20 % of contrast weights with the frontoparietal community in each of the three partitions. Higher Sørensen-Dice coefficient indicates better correspondence. **c**, Positive task contrast weights within the frontoparietal community in each of the three partitions; adult partition shows significantly stronger positive activation than the clustering partition and WSBM partition (*H* (2) = 740.63, *p* < 2 ×10^−16^). **d**, Overlap of lowest 20 % of contrast weights with the default community in each of the three partitions. Higher Sørensen-Dice coefficient indicates better correspondence. **e**, Negative task contrast weights within the default community in each of the three partitions; clustering partition shows significantly stronger negative deactivation than the adult partition and WSBM partition (*p <* 0.001).

We next sought to investigate whether brain areas that are most strongly involved in working memory are well-captured by the frontoparietal community in our developmental partitions. To do so, we examined the positive contrast weights, quantifying which partition had the strongest activation within the frontoparietal community during the cognitively demanding working memory portion of the task (Figure 6c). We found that the adult partition had the strongest positive activation (*M =* 0.38, *SD =* 0.24), with the clustering partition also showing strongly positive contrast weights (*M =* 0.32, *SD =* 0.20) and the WSBM partition showing the least positive task activation (*M =* 0.27, *SD =* 0.23, Kruskal-Wallis test, *H* (2) = 740.63, *p* < 2 ×10^−16^, all pairwise comparisons were significant, *p* < 2 ×10^−16^). Overall, these analyses of task activity suggest that the adult clustering partition corresponds best to task-evoked activity during the n-back task, though the child clustering partition also corresponds well.

#### Default community

Next, we turned to areas of deactivation during the n-back task, examining whether differences in the topography of the default community found in our developmental partitions were reflected in the spatial extent of deactivation during the n-back task. We thresholded the task activation maps to retain only the lowest 20 % of contrast weights (Figure 6b), and compared these to the default communities in each partition. The clustering partition shows the best correspondence to deactivation in the n-back task (Sørensen-Dice coefficient= 0.7273 ±0.001(0.7248 − 0.7299)), with the adult partition also showing correspondence (Sørensen-Dice coefficient = 0.6779 ±0.001(0.6753 − 0.6802)) and the WSBM partition shows the worst performance (Sørensen-Dice coefficient = 0.6496 ± 0.001(0.6473 − 0.6518)). This set of findings suggests that the clustering partition captures community structure that corresponds best to deactivation during tasks.

We next investigated whether brain areas that are deactivated during the cognitively-undemanding portion of the n-back task are well-captured by the default community in our developmental partitions. We examined the negative contrast weights within the default community of each partition, to quantify which partition had the strongest deactivation during the cognitively undemanding 0-back portion of the task (Figure 6d). We found that again, the clustering partition had the strongest deactivation (*M =* −0.23, *SD =* 0.19), with the adult partition also showing strongly negative contrast weights (*M =* −0.20, *SD =* 0.14) and the WSBM partition showing the least negative task deactivation (*M =* −0.17, *SD =* 0.12, Kruskal-Wallis test, (*H* (2) *=* 296.67, *p <* 2 *×* 10^−16^), all pairwise comparisons significant, *p <* 0.001). This set of findings again suggests that the clustering partition captures community structure that corresponds best to deactivation during tasks.

## DISCUSSION

Does the architecture of children’s functional brain networks differ from that of adults? Using a data-driven community detection approach, we found that sensory and motor communities resembled those of adults, but the limbic community was expanded into areas typically assigned the default system in adults, and ventral somatomotor areas were assigned to the ventral attention community and clustered with language-related brain regions. To further probe the interactions between communities, we turned to a model-based approach called the weighted stochastic block model (WSBM). We found a diffuse, scattered pattern of community assignments in prefrontal cortex, perhaps indicative of broadly less-solidified higher-order community interactions and boundaries in children of this age, and again found limbic representation in lateral prefrontal cortex not seen in adults. Across both approaches, the greatest uncertainty in algorithmic assignment of regions to communities was localized to the dorsal and ventral attention communities, including cingulo-opercular regions. Replication in another dataset yielded consistent community assignments for both methods. Activation and deactivation patterns during a working memory task showed that the clustering partition, and the adult partition, captured functional organization in middle childhood well.

The relative expansion of limbic spatial territory in the developmental clustering partition compared to the adult partition is suggestive of a higher importance of limbic circuitry in children^48^, and potentially consistent with compression of limbic territory as emotion regulation abilities develop^16,49^. It is worth noting that while we focus on the adult partition estimated by Ref.^28^, many if not all adult partitions display a hub of the default community in medial prefrontal cortex (e.g. Refs.^3,50^). We also observe some limbic community representation in lateral prefrontal cortex in the developmental WSBM partition, which is not typically observed in adult partitions.

### Variation in transmodal association cortex

We find that community assignments in higher-order association cortex differ the most across approaches. This pattern is consistent with the slow structural development of association cortex, reflected in a prolonged course of thinning and myelination^20,21,23,51^, and with previous findings in children of this age, showing similar community boundaries in primary sensory areas as adults^25^. It is also consistent with the finding that, through adolescence and into adulthood, community assignment remains most variable across individuals in association cortex^2,27,52^. In prefrontal regions of the child clustering partition, we observe substantial changes in assignment between children and adults in frontoparietal, default, and limbic communities, perhaps indicative of unsolidified community assignments in these regions, consistent with evidence that in adults interindividual variability is highest in these areas^6,8^. Another indication of increased flexibility in these areas is the consistency of community assignments across datasets: while the clustering approach yields an almost identical partition in a replication dataset, the WSBM approach shows some differences in assignment, primarily in prefrontal cortex, in the limbic, frontoparietal, and attentional communities.

Even within association cortex, we observe variability in the extent to which children’s assignments look like adults. We see gradients from low to high consistency along the posterior-anterior and medial-lateral axes, with posterior medial regions showing the most consistent community assignments across partitions. In both developmental partitions, the dorsal attention community aligns well with the adult definition, and with known anatomy, encompassing the frontal eye fields and superior parietal lobule^10^. The posterior midline hubs of the default community, the precuneus and posterior cingulate, maintain similar community assignment across the adult partition and both developmental partitions. This posterior hub of the default community plays a mature role even in infant brain networks^53^, and thus its adult-like community assignment in middle childhood and its strong deactivation during the working memory task may be unsurprising evidence for maturity. The evidence for earlier maturity of posterior default hubs, relative to anterior hubs, is mixed, however, as children also show lower confidence in the precuneus and posterior cingulate than adults. In the developmental clustering partition, the ventral aspects of the somatomotor community, which encode the tongue and head^44^, are clustered with the ventral attention community and language-related brain regions, suggestive that, in children, language processes may be more integrated with motor regions involved in the production of language as expertise continues to be built^54^. Similarly, fractionation of the somatomotor community into a set of regions encoding the tongue and head clustered with parts of auditory cortex, was previously observed in the 17-community solution in the adult clustering partition derived by Yeo *et al*.^28^. In the WSBM partition, the ventral aspects of the somatomotor community are but part of an expanded somatomotor community, which extends again to cover language-related brain regions, covering similar territory to the ventral attention community in the clustering partition. It is also possible that the topographic arrangement of the community boundaries in this area of the brain arises simply from the proximity of these regions in volumetric space, as there is some possibility of blurring across sulci.

We focused specifically on localizing the frontoparietal and default communities when probing the relationship between children’s community topography and their patterns of task activity, as we were constrained to the limited set of tasks collected in the full ABCD sample^47^. Despite the variation in community assignment we observed in transmodal association cortex, it seems that both the adult partition by Ref.^28^ and the developmental clustering partition align well with patterns of functional organization as indicated by task activity. This is somewhat unsurprising, as the default community in particular shows some evidence for maturity in posterior areas, and has been shown to be present even in infancy^53^, and the posterior hubs of the default community are consistent across the adult and child clustering partitions. The poor performance of the WSBM partition, however, is particularly notable when examining the frontoparietal community results: there is a clear lack of overlap between the frontoparietal community in the WSBM partition and the areas of high task activation. These results suggest that the flexible interactions between higher-order communities at rest–as implied by the diffuse, scattered community assignments in lateral prefrontal cortex–may “firm up” in the context of a cognitively demanding task, when children shows patterns of functional brain activity that are more adult-like; prior work has shown that functional community organization reconfigures during task performance^55^. To stringently test these partitions, however, ideally we would use several different tasks that tapped the cognitive functions subserved by several sets of communities that show differences in assignment *(e*.*g*. limbic, ventral attention), and this was not possible within the scope of tasks collected in the ABCD sample.

### Algorithmic uncertainty in community assignment

Across both partitioning approaches, areas of highest uncertainty are located in dorsal and ventral attention communities. In the developmental clustering partition, regions assigned to the dorsal attention community show the highest uncertainty, while in the WSBM partition, regions assigned to the ventral attention community show the highest uncertainty, indicative of still-maturing attentional processes in middle childhood^17,18^. Specifically, the anterior insula and anterior cingulate cortex, core components of the midcingulo-insular or “salience” community^3^, show high uncertainty across both approaches. These brain areas are involved in the detection of relevant environmental stimuli (hence the term “salience”) and flexible switching between other large-scale communities^56,57^. Recent work in adults demonstrates that these regions show altered connectivity in response to recent experience^58^, suggesting that they play a key role in flexibly modulating communication with other large-scale communities even in adulthood.

High uncertainty in attention communities may also reflect developing interactions with visual regions. In the developmental clustering partition, the areas of overall lowest confidence are along the border between the visual and dorsal attention communities. This pattern suggests the protracted development of higher-order visual cortex in relation to dorsal attentional circuitry, and is consistent with the slight increase in surface area allocated to the visual community in the developmental clustering partition compared to the adult clustering partition. We observe the assignment of a coherent region in the superior parietal lobule, adjacent to the intraparietal sulcus, to the visual community in the child clustering partition. This area is typically involved in perception of space, spatially-coordinated movements, and magnitude^59^. In the adult partition, a smaller region in the superior parietal lobule is assigned to the visual system, suggesting that an expanded part of this area may be more tightly linked to extrastriate cortex (area MT and anterior MT) in middle childhood^28^.

### Methodological considerations

We first employed a well-established data-driven clustering approach developed by Yeo *et al*.^28^, then used the model-based generative WSBM^40^ to further probe interactions between communities. The two approaches are both similar and distinct. They are similar in that they both attempt to group regions with similar brain-wide patterns of functional connectivity into communities. The WSBM in particular can capture both modular and non-modular types of community structure, with its ability to detect a diverse set of network architectures. The WSBM bears more resemblance to the clustering algorithm than do methods like modularity maximization, which simply seek to maximize connectivity within communities, irrespective of similarities and differences in regional connections between communities. However, the two approaches also have important differences. The WSBM has a precise motivating approach, assuming that nodes can be partitioned such that the distribution of edge weights between sets of communities is governed by parameterized generative processes, and that the parameterization of these processes depends only on the communities to which nodes are assigned. The clustering algorithm, on the other hand, uses a mixture model to cluster regions in a complex high-dimensional space.

We followed Ref.^28^ for the clustering approach and thresholded subject connectivity profiles at 10 % density (i.e., retained only the most positive 10 % of connections), while in the WSBM we were able to retain all edge weights, both negative and positive. This difference implies that the clustering method may be more reliant on strong edges that occur with high frequency across subjects, while the WSBM explicitly groups all nodes into communities with similar connectivity patterns. If weak or negative edges are more variable or noisier than strong edges, the prevalence of weak edges in the unthresholded WSBM approach may account for the diffuse, scattered pattern of community assignments seen in the child WSBM partition in prefrontal cortex. This possibility is also consistent with the pattern observed in our replication dataset: when using the clustering approach, which relies on only the strongest connections, we find a very consistent pattern of community assignments, but when using the WSBM, which incorporates a range of connection strengths, we find more variability in assignments in lateral prefrontal cortex. This difference between approaches–the clustering approach being driven by strong edges, and the WSBM being driven by all edges–may explain the high reproducibility of the clustering approach, but might also sacrifice sensitivity to individual variation, as a growing literature has suggested that middling strength or weak edges best reflect individual differences in cognition^60,61^.

### Limitations

A few limitations of this study should be noted. Most importantly, we do not know ground truth: we are not able to validate our *in vivo* estimates of functional network community structure with histological pediatric data. Pediatric *ex vivo* data are thankfully scarce, but could be helpful in determining which of our two parcellations is most similar to cytoarchitecture or myeloarchitecture. For example, in the WSBM partition but not in the child clustering partition, we observe a patch of the ventral attention community in middle frontal gyrus that resembles a pattern seen in the adult partition. Histology work in adults confirms the presence of a patch of cortex that is cytoarchitectonically differentiated from the surrounding areas, with an expanded layer IV^62^. Relatedly, we cannot use localizer tasks to determine which partition better matches task activation for each community because the ABCD study only includes a few tasks^63^. Second, motion artifact remains a challenging confound in studies of neurodevelopment. In addition to rigorous quality assurance protocols and validated image preprocessing to reduce the impact of motion, we deliberately examined only a subsample of low-motion participants. While this approach mitigates the possible impact of motion artifact on our findings, it may have reduced the generalizability of our sample. Third, imaging techniques have progressed considerably in the last decade, and there are differences in scan acquisition parameters between the dataset used by Ref.^28^ to estimate the adult clustering partition and the child dataset we employ here. This may have led to differences in the distribution of signal to noise ratio (SNR) across the brain. There have been concerns that low SNR in orbitofrontal cortex and temporal pole might hamper community definition in limbic cortex in adults^64^. We conducted additional analyses to rule out the possibility that differences in SNR fully account for the communities we detect in children. Finally, due to computational limitations of the model-based approach, we took a dimensionality-reduction step prior to employing the WSBM, and used a parcellation to downsample the data, rather than using full vertex-wise data. This was a necessary step to make the computations of the WSBM tractable; however, true community boundaries might not be captured if they cut through parcels.

### Broader implications and future directions

Our work builds on prior literature in adults showing that cortex can be divided into reproducible group-average functional communities^3,28,50^. Studies examining functional brain networks typically must choose a partition to apply to characterize their results at the mesoscale or regional level, and we show that a commonly used adult partition may not accurately capture the community structure of children’s brain networks. Instead, we generate two new developmental partitions that can be used for future analyses examining functional brain networks in children of this age. While there is a growing movement towards individualized functional communities, group-level partitions are still widely used, as they allow researchers to easily compare results across participants without the confound of differences in the size or number of communities. Further, individualized approaches typically require large amounts of data per individual^9,65,66^, and thus may not always be feasible in developmental studies.

Many important questions remain. First, how does the architecture of children’s brain networks change as they mature? Future work with longitudinal data will allow us to examine the trajectory of developmental change during the dynamic period of adolescence, and to track how community boundaries shift and stabilize during this time period. Second, what is the cognitive significance of different patterns of community structure in childhood? Answering this question will require well-designed measures of cognition (see Ref.^67^ for an examination of the reliability of the main cognitive assessment tool in the ABCD study). Recent work with the ABCD sample has revealed that the true effect sizes of brain-cognition relationships in this sample are smaller than would be expected based on prior literature^68^, but some studies have shown relationships between variation in functional community topography and cognition^27,52^. Finally, are children’s brains simply less well-captured by partitions into separate communities than adults brains? A group-level partition may not be as useful in children as in adults if children’s brains are simply more variable than adults, and if some regions of cortex are not well-captured by a single community assignment. Our examination of uncertainty in community assignment suggests that capturing this variability is important, and that some brain regions are still highly flexible in middle childhood. Prioritizing methods that estimate the uncertainty and variation in community assignment, such as soft partitioning approaches that assign weighted probabilities of community assignment to each vertex (e.g., Refs.^27,66^), will enable us to test whether functional brain network architecture becomes more solidified as children grow up.

In sum, this work emphasizes the utility of approaches that capture variability and uncertainty in brain network organization. Our findings suggest that one key developmental process might be increasing solidity of brain network architecture as children develop, and set the stage for both theory change and insight into the protracted period of human childhood. These results advance our knowledge about the organization of children’s brains, suggesting a greater representation of regions involved in emotion processing, greater integration of language and somatomotor systems, and more uncertainty in the assignment of association cortex to communities relative to adults. These findings broadly align with differences in the mental lives of children and adults, and with theories about enhanced plasticity in childhood.

## ACKNOWLEDGEMENTS

The authors would like to acknowledge Dr. Richard Watts, who provided the task activation maps used in this study, and thank Dr. Linden Parkes for his helpful comments on an earlier version of this manuscript. U.A.T. was supported by the National Science Foundation Graduate Research Fellowship. A.P.M. was supported by a Jacobs Foundation Early Career Research Fellowship and the National Institute on Drug Abuse (1R34DA050297-01). D.S.B. was supported by the Army Research Office (Grafton-W911NF-16-1-0474) and the National Science Foundation (NSF PHY-1554488, BCS-1631550, and IIS-1926757). Data used in the preparation of this article were obtained from the Adolescent Brain Cognitive Development (ABCD) Study (https://abcdstudy.org), held in the NIMH Data Archive (NDA). This is a multisite, longitudinal study designed to recruit more than 10,000 children age 9-10 years and follow them over 10 years into early adulthood. The ABCD Study is supported by the National Institutes of Health and additional federal partners under award numbers U01DA041048, U01DA050989, U01DA051016, U01DA041022, U01DA051018, U01DA051037, U01DA050987, U01DA041174, U01DA041106, U01DA041117, U01DA041028, U01DA041134, U01DA050988, U01DA051039, U01DA041156, U01DA041025, U01DA041120, U01DA051038, U01DA041148, U01DA041093, U01DA041089, U24DA041123, U24DA041147. A full list of supporters is available at https://abcdstudy.org/nih-collaborators. A listing of participating sites and a complete listing of the study investigators can be found at https://abcdstudy.org/principal-investigators.html. ABCD consortium investigators designed and implemented the study and/or provided data but did not participate in analysis or writing of this report. This manuscript reflects the views of the authors and may not reflect the opinions or views of the NIH or ABCD consortium investigators.

The ABCD data repository grows and changes over time. The ABCD data used in this report came from the Fast Track data release. The raw data are available at https://nda.nih.gov/edit_collection.html?id=2573. Instructions on how to create a NDA study are available at https://data-archive.nimh.nih.gov/training/modules/study.html.

## AUTHOR CONTRIBUTIONS

Conceptualization, U.A.T., D.S.B., and A.P.M.; Methodology, U.A.T., D.S.B., and A.P.M.; Formal analysis, U.A.T.; Writing – Original Draft, U.A.T. and A.P.M.; Writing – Review & Editing, U.A.T., D.S.B., and A.P.M.; Resources, A.P.M.; Supervision, D.S.B. and A.P.M.

## DECLARATION OF INTERESTS

The authors declare no competing interests.

## CITATION DIVERSITY STATEMENT

Recent work in neuroscience and other fields has identified a bias in citation practices such that papers from women and other minorities are under-cited relative to the number of such papers in the field^69–71^. We obtained predicted gender of the first and last author of each reference by using databases that store the probability of a name being carried by a woman^72,73^. By this measure (and excluding self-citations to the authors of our current paper), our references contain 18 % woman(first author)/woman(last author), 9 % man/woman, 27 % woman/man, and 46 % man/man. This method is limited in that a) names, pronouns, and social media profiles used to construct the databases may not, in every case, be indicative of gender identity and b) it cannot account for intersex, non-binary, or transgender people. We look forward to future work that could help us to better understand how to support equitable practices in science.

## STAR METHODS

### Resource Availability

#### Lead Contact

Further information and requests for resources and reagents should be directed to and will be fulfilled by the Lead Contact, Allyson Mackey.

#### Materials Availability

We provide two freely available partitions (in fsaverage6, fsLR, and MNI volumetric spaces), at https://github.com/utooley/Tooley_2020_child_functional_comms/tree/master/partitions.

#### Data and Code Availability

The ABCD dataset (https://abcdstudy.org) is freely available from the NIMH Data Archive (NDA). The ABCD data used in this report came from the Fast Track data release. The raw data are available at https://nda.nih.gov/edit_collection.html?id=2573.

All other analysis code is available at https://github.com/utooley/Tooley_2020_child_functional_comms, along with the two developmental partitions generated in this study. Other toolboxes used in this project are available at https://github.com/ThomasYeoLab/CBIG/tree/master/stable_projects and https://aaronclauset.github.io/wsbm.

### Experimental Model and Subject Details

#### Human participants

Data are from 670 children in the Adolescent Brain Cognitive Development (ABCD) study at the first time point (Release 2.0.1), between the ages of 9 and 11 years (*M =* 9.94, *SD =* 0.67, 47.61% female)^74^. Children were recruited from schools at 21 sites across the United States of America. Due to known scanner effects in the ABCD study^25,63,75^, we selected a subset of children from one scanner at one site who had at least one usable T1-weighted (T1w) image (passed ABCD’s Freesurfer visual assessment checks), had 2 or more resting-state functional magnetic resonance imaging (fMRI) runs with average framewise displacement < 0.5 mm and < 50% of volumes > 0.2 mm^76^, and had an average framewise displacement over all runs < 0.2 mm, after correcting for respiratory artifacts^77^. These parameters were chosen to ensure that we could retain as many participants with high-quality resting-state data as possible, while minimizing the effect of motion on our analyses. Participants were 84 % white, 9 % Hispanic, 6 % other, *<* 1 % Black, and < 1 % Asian. Average parental education ranged from 7 to 20 years (*M =* 15.41, *SD =* 1.88 years, 52 % with bachelor’s degree or higher education).

Replication data are from 544 children from two ABCD sites between the ages of 9 and 11 years (*M* 1=0.2, *SD=*0.53, 55.7% female). Both sites used Siemens scanners. All replication sample participants met the above imaging data quality criteria. Replication sample participants were 74 % white, 15 % Black, 8 % other, 3 % Hispanic, and < 1 % Asian. Average parental education ranged from 10 to 20 years (*M =*15.36, *SD =*2 years, 48 % with bachelor’s degree or higher education).

### Method Details

#### Image Acquisition

The ABCD scan session included T1w and T2-weighted (T2w) images, one dMRI series, four 5-minute resting-state fMRI series, and three sets of two task fMRI series. One set of two 5-min resting-state fMRI runs is acquired immediately after the T1w scan and another set is acquired after the T2w scans, followed by task fMRI runs. Resting-state data were acquired with eyes open during passive viewing of a cross hair. The T1w acquisition (1 mm isotropic) is a 3D T1w inversion prepared RF-spoiled gradient echo scan using prospective motion correction^78^. The fMRI acquisitions (2.4 mm isotropic, TR=800 ms) use multiband echo-planar imaging with slice acceleration factor 6. Details about ABCD image acquisition are available elsewhere^63,79^.

#### Data quality and exclusion criteria

Due the nature of the ABCD Fast Track data, which enables almost immediate access to the raw images from this study, there are occasional errors in data quality or subject ID assignment. We excluded any participants from the target site whose data on Amazon S3 was incomplete as of Spring 2019 or whose data contained sequences not officially part of the ABCD study. Runs of resting-state fMRI that contained < 360 volumes or where quality metrics indicated poor coregistration (cross-correlation or Jaccard coefficient < 0.95, mincost of `bbregister` > 0.6) were excluded. Following preprocessing, runs with > 50% of volumes flagged as outliers for average framewise displacement > 0.2 mm were removed from analyses. Any subjects with less than 2 runs of resting-state fMRI data remaining after these exclusions were not included in analyses.

#### Preprocessing

Results included in this manuscript come from preprocessed data, where the preprocessing was performed using *fMRIPprep* 1.4.1 (^80^;^81^; RRID:SCR_016216), which is based on *Nipype* 1.2.0(^82^;^83^; RRID:SCR_002502), as well as XCPEngine 1.0^84^.

The T1-weighted (T1w) image was corrected for intensity non-uniformity with `N4BiasFieldCorrection`^85^, distributed with ANTs 2.2.0^86^ (RRID:SCR_004757), and used as T1w-reference throughout the workflow. The T1w-reference was then skull-stripped with a *Nipype* implementation of the `antsBrainExtraction.sh` workflow (from ANTs), using OASIS30ANTs as the target template. Brain tissue segmentation of cerebrospinal fluid (CSF), white-matter (WM) and gray-matter (GM) was performed on the brain-extracted T1w using `fast`^87^. Brain surfaces were reconstructed using `recon-all`^88^, and the brain mask estimated previously was refined with a custom variation of the method to reconcile ANTs-derived and FreeSurfer-derived segmentations of the cortical gray-matter of Mindboggle^89^.

For each of the up to 4 resting-state blood oxygen level-dependent (BOLD) runs found per subject (across all tasks and sessions), the following preprocessing was performed. First, a reference volume and its skull-stripped version were generated using a custom methodology of *fMRIPrep*. A deformation field to correct for susceptibility distortions was estimated based on two echo-planar imaging references with opposing phase-encoding directions, using `3dQwarp`^90^ (AFNI 20160207). Based on the estimated susceptibility distortion, an unwarped BOLD reference was calculated for a more accurate co-registration with the anatomical reference. The BOLD reference was then co-registered to the T1w reference using `bbregister` (FreeSurfer) which implements boundary-based registration^91^. Co-registration was configured with nine degrees of freedom to account for distortions remaining in the BOLD reference. Head-motion parameters with respect to the BOLD reference (transformation matrices, and six corresponding rotation and translation parameters) are estimated before any spatiotemporal filtering using `mcflirt`^92^. The BOLD time-series were resampled onto their original, native space by applying a single, composite transform to correct for head-motion and susceptibility distortions. These resampled BOLD time-series will be referred to as preprocessed BOLD in the original space, or just preprocessed BOLD.

Several confounding time-series were calculated based on the preprocessed BOLD: framewise displacement (FD), the rate of change of BOLD signal across the brain at each frame (DVARS), and three region-wise global signals. FD and DVARS are calculated for each functional run, using the implementations in *Nipype*^93^. The three global signals are extracted within the CSF, the WM, and the whole-brain masks. The head-motion estimates calculated in the correction step were also placed within the corresponding confounds file. The confound time series derived from head motion estimates and global signals were expanded with the inclusion of temporal derivatives and quadratic terms for each^94^.

All resamplings can be performed with a single interpolation step by composing all the pertinent transformations (i.e. head-motion transform matrices, susceptibility distortion correction when available, and co-registrations to anatomical spaces). Gridded (volumetric) resamplings were performed using `antsApplyTransforms` (ANTs), configured with Lanczos interpolation to minimize the smoothing effects of other kernels^95^.

Many internal operations of *fMRIPrep* use *Nilearn* 0.5.2^96^ (RRID:SCR_001362), mostly within the functional processing workflow. For more details of the pipeline, see the section corresponding to workflows in *fMRIPrep*’s documentation.

Further preprocessing was performed using a confound regression procedure that has been optimized to reduce the influence of subject motion^94,97^; preprocessing was implemented in XCPEngine 1.0^84^, a multi-modal toolkit that deploys processing instruments from frequently used software libraries, including FSL^98^ and AFNI^99^. Further documentation is available at https://xcpengine.readthedocs.io and https://github.com/PennBBL/xcpEngine. Functional timeseries were band-pass filtered to retain frequencies between 0.01 Hz and 0.08 Hz. Data were demeaned, and linear and quadratic trends were removed. Confound regression was performed using a 36-parameter model; confounds included mean signal from the whole brain, white matter, and CSF compartments, 6 motion parameters as well as their temporal derivatives, quadratic terms, and the temporal derivatives of the quadratic terms^94^. Prior to confound regression, all confound parameters were band-pass filtered in a fashion identical to that applied to the original timeseries data, ensuring comparability of the signals in frequency content^100^. Motion censoring was applied by removing frames with FD > 0.2 mm or standardized DVARS > 2. To avoid variability in scan duration influencing results, the first 2 runs from each subject that met all inclusion criteria were used as input to both partitioning algorithms.

#### Data-driven community detection approach: Clustering algorithm

For both partitions, we first systematically investigated the number of communities, or the optimal *k*, that best describes cortical organization in childhood (see *Number of communities* sections below). Based on our findings, and prior evidence that functional brain networks can be divided into 6-10 communities^28,101–103^, we chose *k* = 7 for our main analyses to facilitate comparison with the adult partition derived in Ref.^28^. The clustering algorithm and the WSBM both attempt to group vertices or parcels into communities based on their patterns of connectivity with the rest of the brain, but the extent of their mathematical definition and the level at which they operate varies.

##### Surface-based processing

Following temporal filtering and confound regression, functional data were projected onto the surface using *mri_vol2surf* (FreeSurfer) and downsampled to fsaverage6 surface space (where the vertex spacing is roughly 2 mm). Data were smoothed using a 4.8 mm full-width half-maximum (FWHM) kernel, similarly to the adult dataset from which the adult partition was derived^28^.

##### Number of communities

We systematically varied the number of communities detected, from *k =*2 − 17, and implemented the vertex-wise instability analysis from Ref.^28^ to examine the optimal *k* for the child clustering partition. Briefly, the instability analysis involves repeatedly and randomly dividing the 74,846 vertices into two groups, and applying the clustering algorithm to each group separately. The parameters learned from clustering the first group of vertices are then used to predict the clustering results for the second set of vertices, and the agreement between the prediction and clustering results of the second group of vertices measures the generalizability of the clustering results at that *k*. The vertex resampling was iterated 100 times at each *k*, with a different random split of vertices into groups each time. All other clustering parameters were set the same as in the clustering algorithm, above. Less stability is observed with increasing number of estimated communities, which is an expected property, since the number of estimated communities enlarges the solution space of the clustering problem. Lower values of instability indicate higher consistency across re-samplings at a given *k*, and thus better partition fit.

##### Clustering algorithm

The clustering algorithm attempts to detect functionally coupled regions and was implemented following Ref.^28^. Here we provide a brief overview of the algorithm for clarity. Time-varying BOLD signals were extracted from each vertex on the surface. Connectivity between each vertex and 1,175 evenly-spaced regions of interest (ROIs) was estimated by calculating the Pearson correlation coefficient between their timeseries and normalized. The resulting 74,846 × 1,175 correlation matrix was averaged across runs, and then binarized to keep the top 10% of the correlations; the resulting connectivity profiles were averaged across subjects. The averaged connectivity profiles were clustered using a mixture of von Mises–Fisher distributions^104^ with *k =* 7 based on our previous results and for ease of comparison with the adult partition. This approach modeled the 74,846 points of data on a 1,174-dimensional hyper-sphere in a 1,175-dimensional space, and attempted to minimize the geodesic distances between points (i.e. attempted to group vertices with similar connectivity profiles to the 1,174 ROIs together in the same community). This procedure also means that vertices were clustered based on their connectivity profiles rather than their absolute connectivity strength; at each iteration the algorithm attempted to maximize the agreement of connectivity profiles within a community. The algorithm was iterated 1000 times with a different random initialization of vertices to communities each time, then the best solution of those 1000 tries chosen based on the likelihood of that partition. The clustering algorithm was implemented using publicly available code from Ref.^28^, using v0.17.0 at https://github.com/ThomasYeoLab/CBIG.

##### Uncertainty in community assignment

As has been done in prior work, we used the silhouette measure^105^, called confidence in Ref.^28^, as a vertex-wise measure of uncertainty in community assignment. The silhouette measure captures the similarity of a vertex’s time-series to other vertices assigned to the same community, compared to the next most similar community. The silhouette for point *i* is defined as:

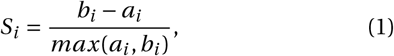

where *a*_*i*_ is the average distance (correlation, in our case) from point *i* to the other points in the same community as *i*, and *b*_*i*_ is the minimum average distance from the *i* th point to points in a different community (minimized over communities). The resulting measure ranges from -1 to 1, with higher values indicating greater confidence in community assignment. Negative values are unlikely, but possible, as the cost function of the clustering algorithm is not equivalent to the silhouette measure. We employ publicly released adult confidence maps from Ref.^28^ for comparison, and as these were estimated in fsaverage5 space, we upsampled them to fsaverage6 space for comparison to the clustering partition.

##### Signal to noise ratio

To estimate the effects of susceptibility artifacts on the child data, we calculated the voxel-wise temporal signal to noise ratio in the child dataset. Following Ref.^28^ and the GSP dataset^64^, we computed the SNR of the fMRI time series for each voxel in subjects’ native volumetric space after pre-processing with *fMRIPprep* by averaging the signal intensity across the whole run and dividing it by the standard deviation over time. SNR maps were averaged across runs, then projected to fsaverage5 surface space and averaged across subjects. Low SNR is present in expected areas, namely, the anterior portion of the inferior and medial temporal lobe, as well as areas of orbitofrontal cortex (see Supplemental Figure S3). We employed publicly released adult SNR maps from Ref.^28^ for comparison with the child SNR data.

#### Model-based community detection approach: Weighted stochastic block model

##### Network construction

Mean BOLD timeseries were extracted from a 400-region parcellation^29^. We estimated the functional connectivity^106^ between any two brain regions by calculating the product-moment correlation coefficient *r*^107^ between the mean activity time series of region *i* and the mean activity time series of region *j* ^108^. Correlations were subsequently *r*-to-*z*-transformed. We represented the *n*× *n* functional connectivity matrix as a graph or network^109^, in which regions were represented by network nodes, and in which the functional connectivity between region *i* and region *j* was represented by the network edge between node *i* and node *j*. We used this encoding of the data as a network to produce an undirected, signed and weighted adjacency matrix **A**. Adjacency matrices were then averaged across the 2 runs, for consistency with the procedures employed in the data-driven approach. We note that despite common use, averaging individual subject matrices to produce a group-average adjacency matrix may result in a structure that is not central to the ensemble of individual matrices^110^.

##### Weighted stochastic block model

Following Ref.^40^, the weighted stochastic block model places each of *n* nodes of the adjacency matrix *A* of subject *f* into one of *k* communities, by maximizing the likelihood that each block of connections between two communities is internally similar. In the classic SBM, the probability of edge existence is learned for each block (system). In the weighted SBM, the edge weight distribution parameterized by *µ* and *σ* is learned for each block. For each subject, we maximize the likelihood of a partition *y* such that *µ* and *σ* parameterize the normally distributed probability of edge weights between nodes in community *i* and community *j*, where nodes in this case are parcels from the Schafer400 parcellation. The WSBM seeks to partition a subject’s brain network such that nodes with similar patterns of connectivity to other nodes are grouped together, under the assumption that each community’s set of edge weights can be modeled with a normal distribution with mu and sigma. This placement is accomplished by finding a network partition *y* ∈*Y* ^*nx*1^ where *y*_*i*_ ∈ 1, 2, …, *k* and *w*_*i*_ denotes the membership of node *i*. Following Refs.^40^ and^42^, we model edge weights with an normal family distribution and discount the contribution of the edge existence distribution. Then the generative model takes the following form:

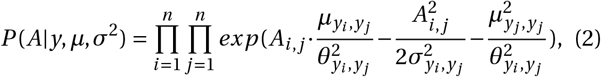

where *µ ∈ R*^*kxk*^ and *σ*^2^ *∈ R*^*kxk*^ are model parameters, and 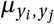 and 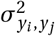 parameterize the weights of normally distributed connections between community *y*_*i*_ and community *y* _*j*_. The quantity *A*_*i, j*_ denotes the *i, j* th element of the network adjacency matrix *A*. The quantity *P* (*A* | *y, µ, σ*^2^) is the probability of generating the observed network *A* given the parameters; this model is fit to *A* to estimate the *y, µ* and *σ*^2^ parameters. For a given subject’s *n* × *n* functional brain network, we maximize the likelihood of the weighted stochastic block model using a variational Bayes technique described by^40^ and implemented in MATLAB code freely available at https://aaronclauset.github.io/wsbm/. We repeated the optimization procedure 50 times for each subject, each time initializing the algorithm with a different set of parameters. We selected *k =* 7 based on our goodness-of-fit results, as well as prior evidence that functional brain networks can be divided into 7 separate components^28^ and to facilitate ease of comparison with our set of *a priori* community assignments. The weighted stochastic block model generated a single maximum likelihood partition of regions into functional communities for each subject.

##### Number of communities

We systematically varied the number of communities detected, from *k=* 2 −17, and implemented analyses of goodness-of-fit to examine the optimal *k* for the child WSBM partition. We first examined the log-likelihood of the WSBM, fit to the main dataset across values of *k* for each subject, repeating the optimization procedure 30 times per subject and each time initializing the algorithm with a different set of parameters. Then, we calculated the log-likelihood of the consensus WSBM partition at a given *k* (derived from our main dataset) fit to the replication dataset. Finally, we examined the number of communities detected at the group level when using the iterative consensus partitioning procedure to derive a representative group WSBM partition at a given *k*.

##### Consensus partition algorithm

To derive a representative consensus WSBM partition, we used an iterative consensus partitioning procedure^111^. This procedure tabulates the co-occurrence of two regions being assigned to the same community across all subjects, subtracts a null model of the chance occurrence of two regions being assigned to the same community, then uses a Louvain-like algorithm to maximize the modularity of the co-occurrence matrix^112^. This final step is iterated *n*=670 times, once for each subject in the sample. We relabeled the communities in the representative consensus WSBM partition using the Hungarian algorithm for maximal overlap with the adult partition communities^113^. Note that although the choice of *k* constrains the number of communities detected at the subject level, we observed that the number of communities detected at the group consensus level can vary from the *k* set at the subject level.

##### Uncertainty in community assignment

As a parcel-wise measure of uncertainty in community assignment, we used the proportion of inconsistent assignments across iterations of our consensus partitioning algorithm. Typically, the consensus partitioning algorithm will consistently assign parcels to the same community across iterations, however, in our case some parcels were inconsistently assigned to communities across iterations. In a supplemental analysis, we also used the silhouette measure introduced above on the WSBM partition, using parcel time-series and WSBM community assignments to estimate the silhouette of each parcel.

#### Partition comparisons

We used information-theoretic measures for clustering comparison to compare the adult partition to the two estimated partitions, namely, the normalized mutual information and the normalized information distance^43^. Permutation tests across vertices (*n =* 81, 924) were conducted to estimate a distribution and calculate a *p*-value. To compare activation and deactivation maps from the n-back task to specific communities, we used measures of set similarity designed for binary comparison, namely the Sørensen-Dice coefficient. All adult partition comparisons were conducted using the publicly released 7-system partition from Ref.^28^ in fsaverage6 space.

#### Uncertainty across both developmental partitions

To examine areas of high uncertainty in assignment across both partitions, we combined the variability in assignment measure from the WSBM with the confidence measure from the clustering algorithm. Percentages of inconsistent assignment in the iterative consensus partitioning procedure were scaled to [0,1] and inverted before summing with confidence maps from the clustering partition. Additionally, we examined at the vertex resolution how many times a vertex changed assignment between the adult partition, the clustering partition, and the WSBM partition (Supplemental Figure S7). This measure ranges simply from 1 to 3, as a vertex can be assigned at maximum to 3 different communities across the 3 partitions.

#### Task-evoked activity

We used maps of task-evoked activity in the emotional n-back task from the ABCD Study^47^. The emotional n-back has both 0-back (low working memory load) and 2-back (high working memory load) conditions. Comparison of the two conditions allows for the assessment of activation specifically related to working memory. Each trial requires a motor response from the subject, specifying whether the stimuli was seen 2 trials ago (in the 2-back condition), or is a target stimuli for the block (0-back). Specifically, we used the 2-back versus 0-back group average-contrast, controlling for age, sex, scanner, and performance on the task.

### Quantification and Statistical Analysis

For information-theoretic measures of partition similarity (normalized mutual information and normalized information distance), we conducted resampling permutation tests to estimate a distribution to calculate a *p*-value. We employed Kruskal-Wallis tests and *post-hoc* Wilcoxon rank sum tests to test differences in uncertainty of community assignments and differences in the contrast weights in our task activation analyses. We bootstrapped standard errors and confidence intervals for binary measures of set similarity (Sørensen-Dice coefficient) using the package *boot* with 1000 repetitions. We conducted all analyses in R and MATLAB using custom code, including that available at https://github.com/ThomasYeoLab/CBIG/tree/master/stable_projects.

#### Data visualization

Surfaces and partitions were shown on cortical surfaces generated by Freesurfer^88^, using *fsbrain* 0.3.0 and *freesurfer-formats* 0.1.11. Connectivity matrices were visualized in MATLAB, all other figures were produced using R^114^.

---

**FIG. S1.**
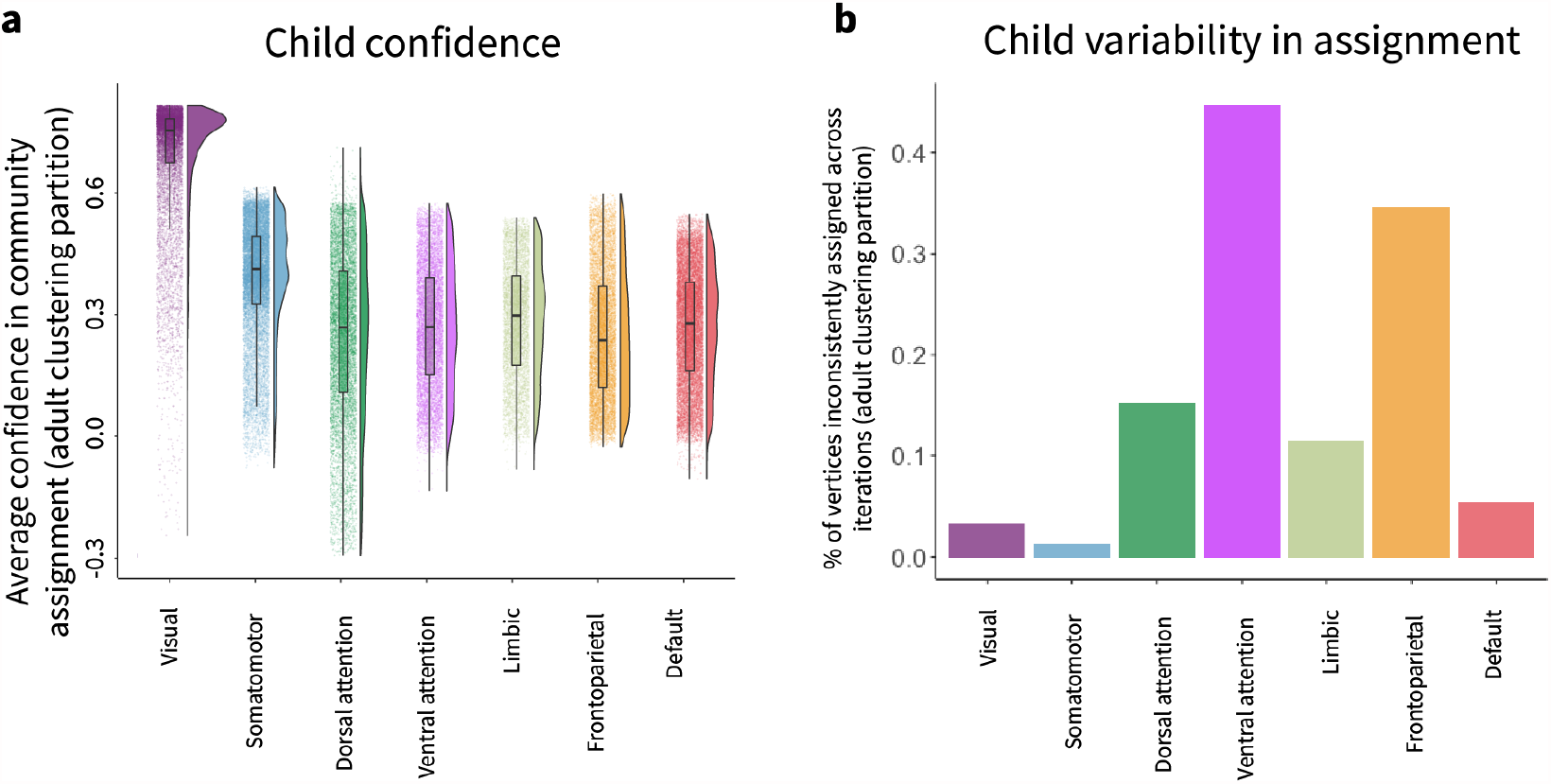
Uncertainty in community assignment using adult clustering partition systems. **a**, Average confidence in the developmental sample, using the data-driven community detection approach, within each of the systems in the adult clustering partition. Note the higher confidence in the visual community and relatively lower confidence in higher-order association systems, particularly the dorsal attention and frontoparietal systems. **b**, Variability in community assignment in the developmental sample, using the model-based community detection approach, within each of the systems in the adult clustering partition. Variability in parcel assignment is predominantly located in the ventral attention and frontoparietal systems.

**FIG. S2.**
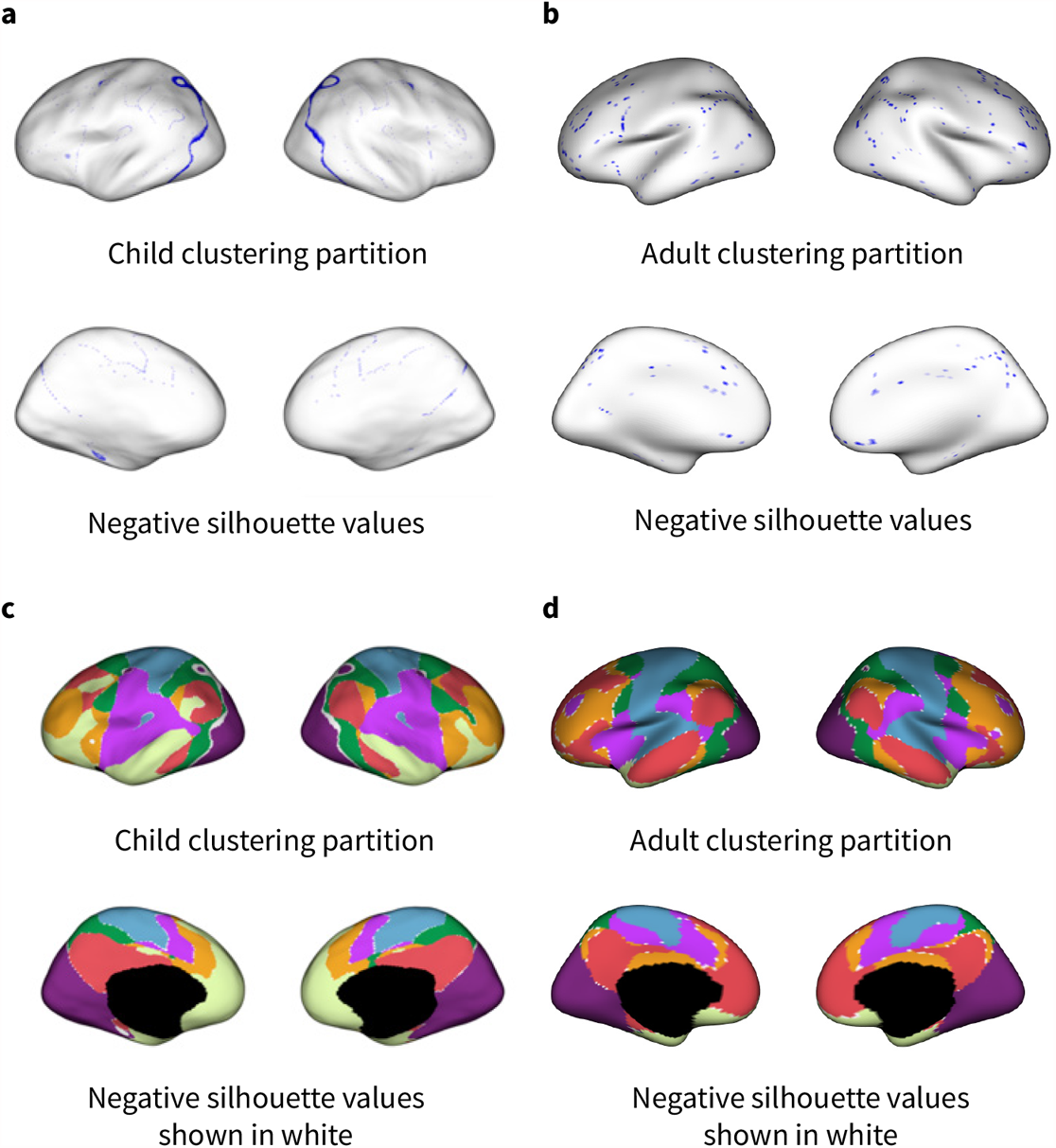
Negative confidence values. **a**, Negative silhouette values in the child clustering partition shown in blue. Negative values are unlikely, but possible, as the cost function of the clustering algorithm is not equivalent to that of the silhouette measure. In the adult sample, negative silhouette values occur along borders between communities (panel **(b)**). Here, too, these values predominantly fall along borders between communities. **b**, Negative silhouette values in the adult clustering partition, data from Ref.^28^. **c**, In the child clustering partition, the vast majority of negative values occur along the border between the visual and dorsal attention communities. These are vertices assigned to the dorsal attention community in the clustering algorithm, but their timeseries are highly similar to that of the visual community. **d**, In the adult clustering partition, negative values occur sporadically along the borders between communities.

**FIG. S3.**
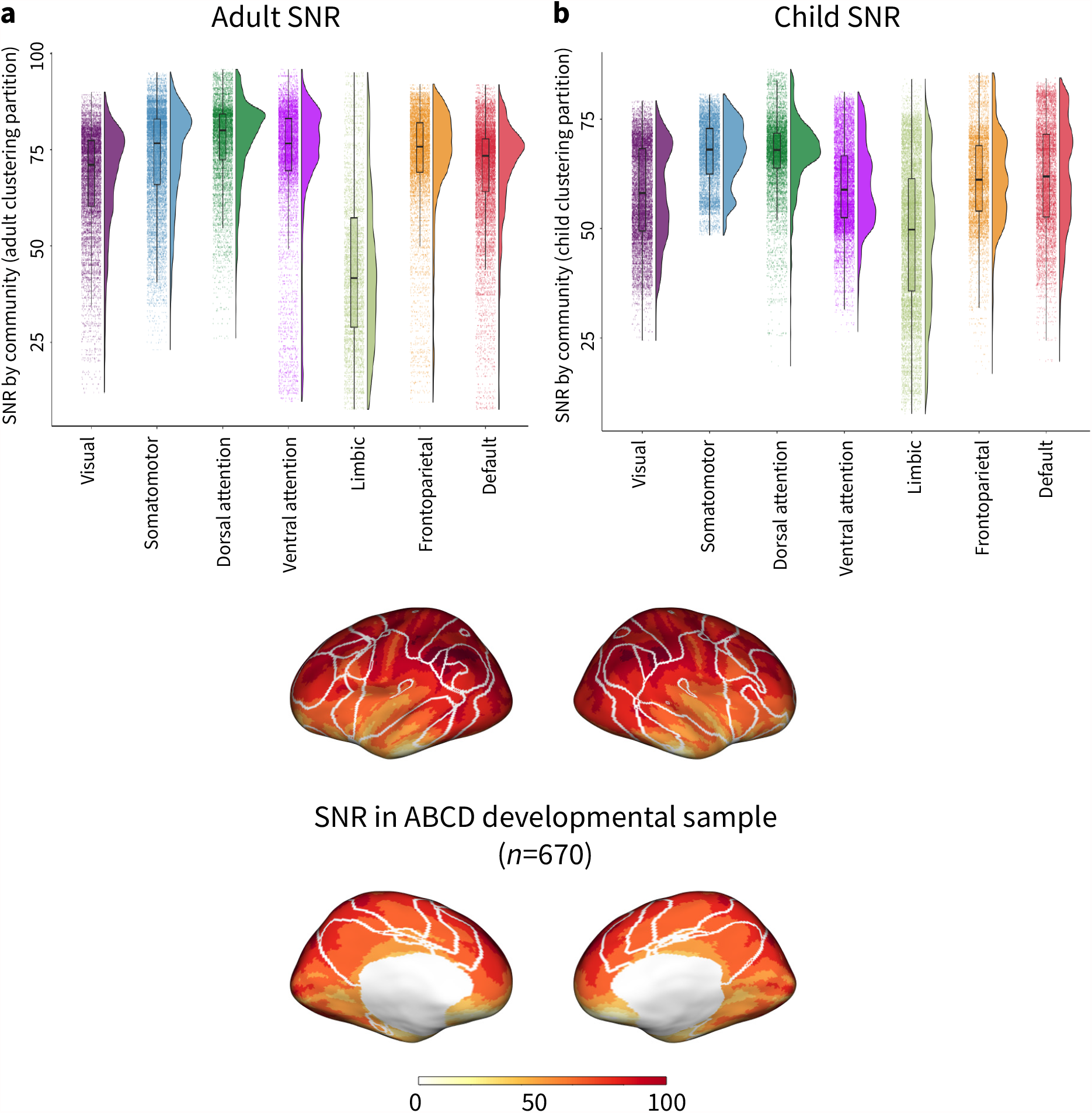
Signal to noise ratio (SNR) in adult and child data. **a**, SNR by adult clustering partition communities, data from Ref.^28^. Variance in SNR in adults is greater within communities than between communities; the limbic community shows the lowest average SNR. **b**, SNR by child clustering partition communities. Though average overall SNR is lower than in the adult data, likely due to differences in scan acquisition parameters between datasets, variation in SNR is again greater within communities than between communities. **c**, In the child clustering partition, the boundaries between communities do not fall neatly along shifts in SNR, indicating that SNR is likely not the primary driver of community assignments.

**FIG. S4.**
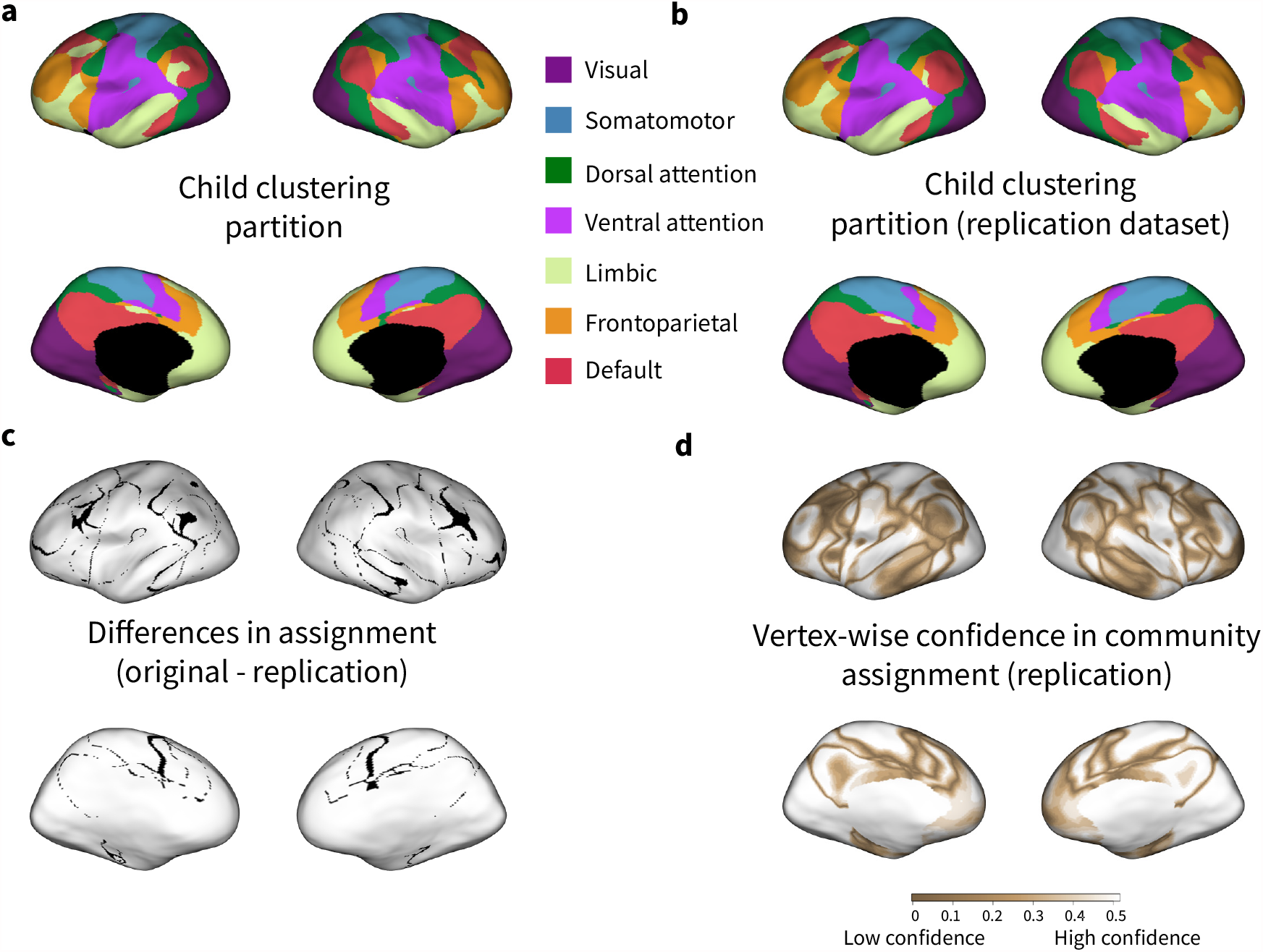
Community assignments using the clustering approach are highly consistent in a replication dataset. **a**, Clustering partition derived from original dataset, shown for visual comparison with panel **(b). b**, Clustering partition derived from the replication dataset. Community assignments are highly consistent across the original and replication datasets; 94.99 % of vertices had the same community assignment across the two datasets. **c**, Vertices with differing community assignments between the original and replication partitions (≈ 5 % of vertices). **d**, Confidence maps from the developmental replication sample, using the silhouette method. Note the visual resemblance to Figure 3b.

**FIG. S5.**
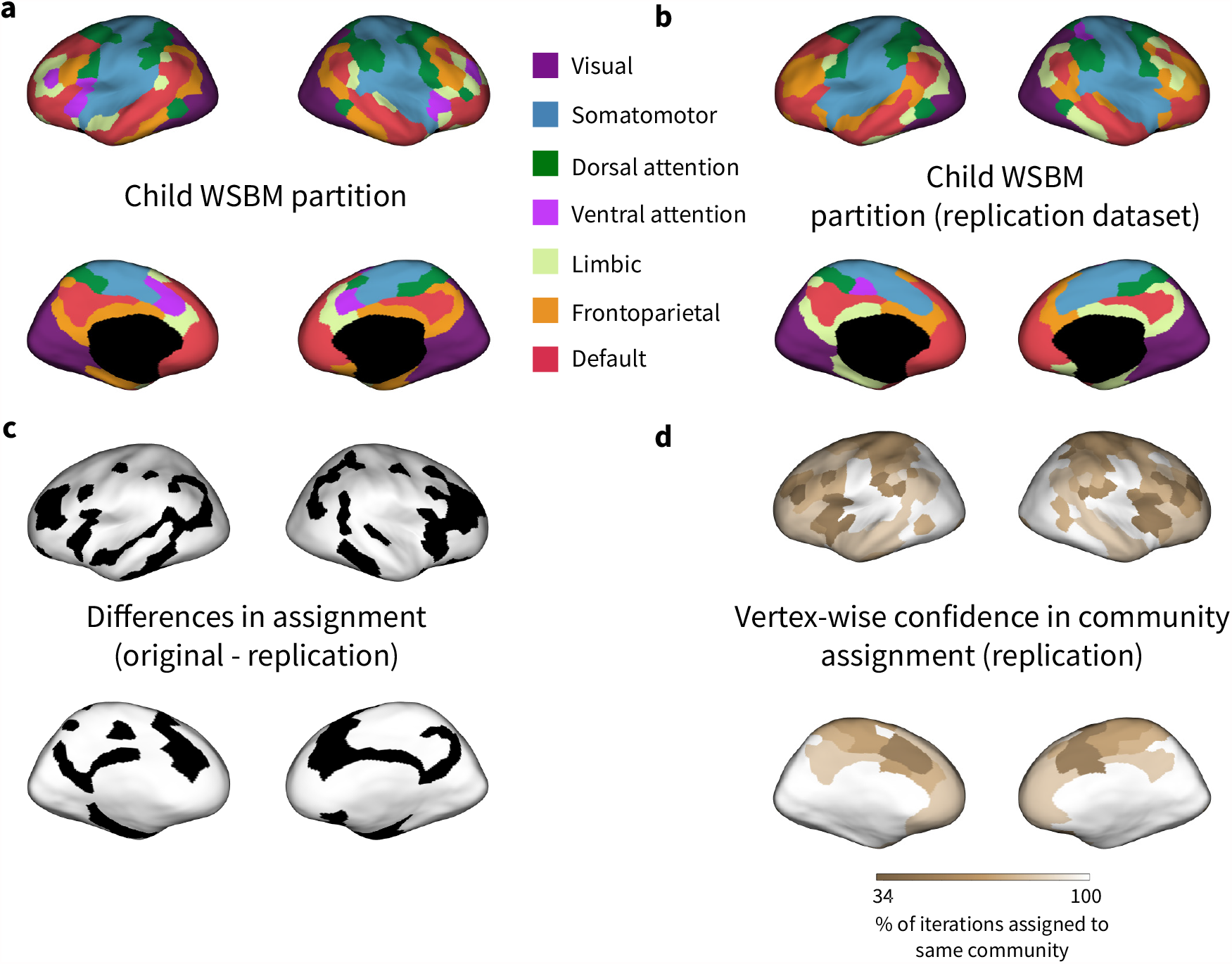
Community assignments using the WSBM approach are highly consistent in a replication dataset. **a**, WSBM partition derived from original dataset, shown for visual comparison with panel b). **b**, WSBM partition derived from replication dataset. Community assignments are consistent across the original and replication datasets; 74.62 % of parcels had the same community assignment across the two datasets. **c**, Parcels with differing community assignments between the original and replication partitions (≈ 15 % of vertices). In keeping with other findings, most variation in community assignment is in lateral prefrontal cortex. **d**, Parcels that were inconsistently assigned to communities during the optimizations of the WSBM consensus partition algorithm.

**FIG. S6.**
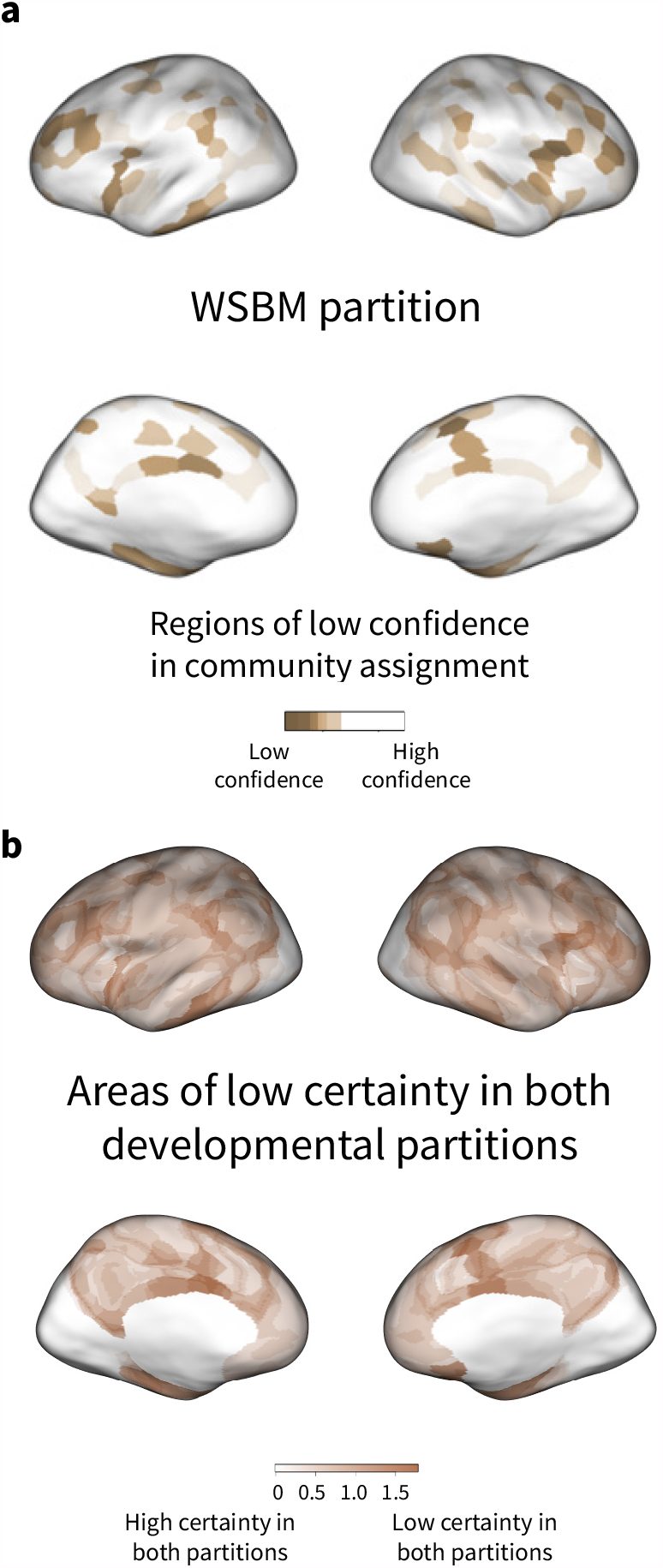
Confidence in community assignment using a model-based (WSBM) approach. **a**, Confidence maps estimated from the developmental sample using the silhouette method. This method measures the similarity of a given vertex’s timeseries to that of other vertices assigned to the same community, compared to the next most similar community (see Methods). For the purposes of visualization, negative silhouette values were set to 0. **c**, Areas of low certainty in community assignments across both developmental partitioning approaches. Regions of low certainty are shown in dark sandstone. Confidence values from the WSBM partition are scaled to [0,1], then summed with confidence values from the clustering partition scaled to [0,1]. Values close to zero show high certainty in both partitioning approaches.

**FIG. S7.**
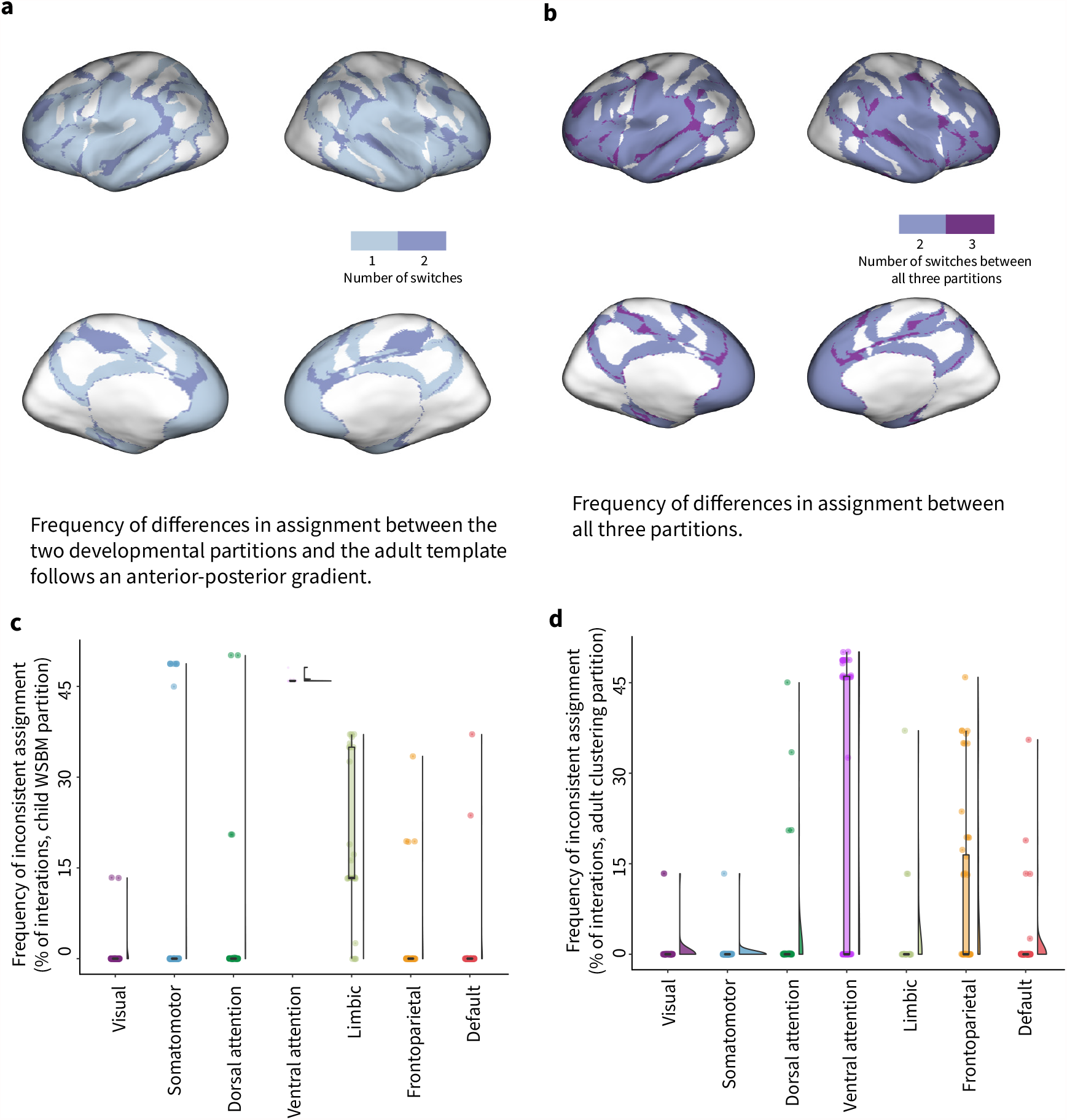
Frequency of changes in community assignment follows an anterior-posterior gradient. **a**, Number of different community assignments for each vertex, calculated between the two developmental partitions and the adult clustering partition. **b**, Number of different community assignments for each vertex, calculated across all three partitions. **c**, Proportions of variability in community assignment in the model-based (WSBM) approach. As an alternative to visualizing the percentage of parcels that had any inconsistent assignments across optimizations of the WSBM consensus community algorithm, we examined the proportions of inconsistent community assignments across optimizations by child WSBM partition communities. Each datapoint is a parcel; 52 % of parcels were consistently assigned to the same community across all optimizations of the consensus partitioning algorithm, resulting in many datapoints at 0. Variability in community assignment is predominantly located in the ventral attention and limbic communities. **d**, Proportions of inconsistent community assignment by adult clustering partition systems. Variability in community assignment is predominantly located in the ventral attention and frontoparietal systems.

## Notes

### Competing Interest Statement

The authors have declared no competing interest.

